# A layer-resolved diagnostic identifies bias-driven decisions in deep neural networks

**DOI:** 10.1101/2025.09.16.676625

**Authors:** Johan Nakuci

## Abstract

Modern AI systems can be accurate and confident, but this alone does not reveal whether a decision is well supported by the input. This creates a trust problem because confidence reports how decisive a model is, but not what supports that decisiveness. Here we show that neural-network confidence can be decomposed into input-dependent feature support and input-independent offset support. We formalize this decomposition through the Bias Dominance Index (BDI), a layer-resolved measure quantifying the relative contribution of input-independent offsets to the decision margin, revealing whether confidence is primarily feature-supported or bias-driven. Across convolutional neural networks, a vision transformer and a transformer language model, BDI shows that high confidence can coexist with bias-driven decisions. Layer-resolved analyses map where bias-driven support across network depth. Perturbation analyses further show that the bias component can stabilize performance when readout weights are degraded. Finally, we operationalize decision composition into an acceptance rule that combines confidence and BDI for mechanism-aware auditing and triage. Together, these results position BDI as a general diagnostic of decision composition that distinguishes feature-supported from bias-driven decisions across model families.

## Introduction

Deep neural networks (DNNs) increasingly operate as decision-making systems in settings where inputs are incomplete, ambiguous, or degraded – images are occluded^1^, sensor streams are noisy^2,3^, and language inputs are underspecified^4,5^ or adversarially framed^3,6^. In these situations, models often remain decisive, assigning high confidence to predictions even when the input provides limited support^7^. This creates a practical trust problem.

For deployment, the central question is not only whether a prediction is confident, but what computational substrate supports that confidence. Confidence is expressed through the decision margin and a large decision margin can arise because the current input provides strong evidence, or because input-independent offsets, such as learned bias terms, normalization shifts, or learned baselines, contribute substantially regardless of the specific input^8^. These offsets are a normal and useful part of neural-network computation, but they matter for trust because they can help sustain decisiveness even when input-dependent evidence is weak. The practical challenge is therefore not simply whether to trust a model globally, but when to trust a particular decision. This problem of trust requires diagnostics that go beyond performance and confidence to ask what supports the model’s decisiveness on each input.

A substantial body of work provides complementary tools for probing these issues. Uncertainty estimation and calibration relate confidence to error^9,10^, and attribution methods map outputs to influential input features or internal units^11–14^. These tools are useful, but they address different aspects of model behavior. Calibration asks whether confidence is statistically warranted, and attribution asks which features influence the output. Neither directly explains what computational components support the model’s confidence on a given input. This distinction matters because a prediction can be both correct and confident while differing in whether its margin is primarily feature-supported or bias-driven. From a trust perspective, confidence alone would treat these cases as equivalent, even though their mechanistic basis differs. As a consequence, confidence can mask distinct forms of decision support.

Importantly, here bias denotes an intercept-like contribution to the logit margin that is independent of the current input. Therefore, the bias in the current analysis is an input-independent offset, rather than bias arising from training-set composition, label imbalance, systematic miscalibration or dataset artifacts.

Addressing this issue requires moving beyond the magnitude of the decision margin to its composition, asking whether the margin is supported primarily by input-dependent features or by input-independent offsets. We formalize this decomposition through the Bias Dominance Index (BDI), a measure that quantifies the relative contribution of input independent offsets (i.e. bias) to the decision margin. For a broad class of neural networks, the decision margin can be decomposed into feature and bias components, enabling BDI to distinguish decisions in which confidence is primarily feature-supported from those in which bias disproportionately shape the output. Thus, BDI is a mechanism-aware diagnostic of margin composition. Crucially, the goal is not merely to restate that bias terms exist, but to use their relative dominance to identify bias-driven decisions and improve trust.

We extend the diagnostic across depth by partitioning each layer’s gradient-weighted local signal into feature and input-independent components. This yields a depth-dependent map of the decision margin, providing a profile of bias dominance across layers of a network. Additionally, we adapt the framework to multi-class models using a label-free decision margin defined by the top-1 versus top-2 logit difference, because deployment settings often lack labels enabling BDI-based auditing of decisions without ground-truth annotations.

First, we apply these diagnostics to a set of convolutional neural networks (CNNs) trained on a controlled perceptual decision-making task. BDI reveals systematic decisions in which feature contributions are attenuated and the bias component disproportionately supports the margin, while layer-resolved BDI maps where this mode across network depth. Additionally, we test the mechanistic relevance of the bias component through perturbations that selectively degrade feature-carrying computation. We then establish that the BDI signature generalizes to other convolutional networks (AlexNet^15^ and VGG-16^16^), a vision transformer (ViT-b/16^17^), and a transformer language model (BERT^18^). Similarly in these networks, BDI flags bias-dominant yet confident decisions, mapping bias dominance in the network hierarchy, and provides a framework for auditing and stress-testing models under degraded feature and distributional uncertainty. Lastly, we operationalize decision composition into a two-parameter acceptance rule that combines confidence with BDI for auditing. Together, these results position BDI as a practical diagnostic for interpretability and reliability in modern neural networks.

### Decision-margin decomposition and the Bias Dominance Index

To formalize how a DNN’s decision is supported, we consider the decision margin between *K* classes. For an input *x*, logits *z*(*x*) are produced by an affine readout from the final internal activation vector *a* (**Fig. 1A, B**):

**Figure 1.**
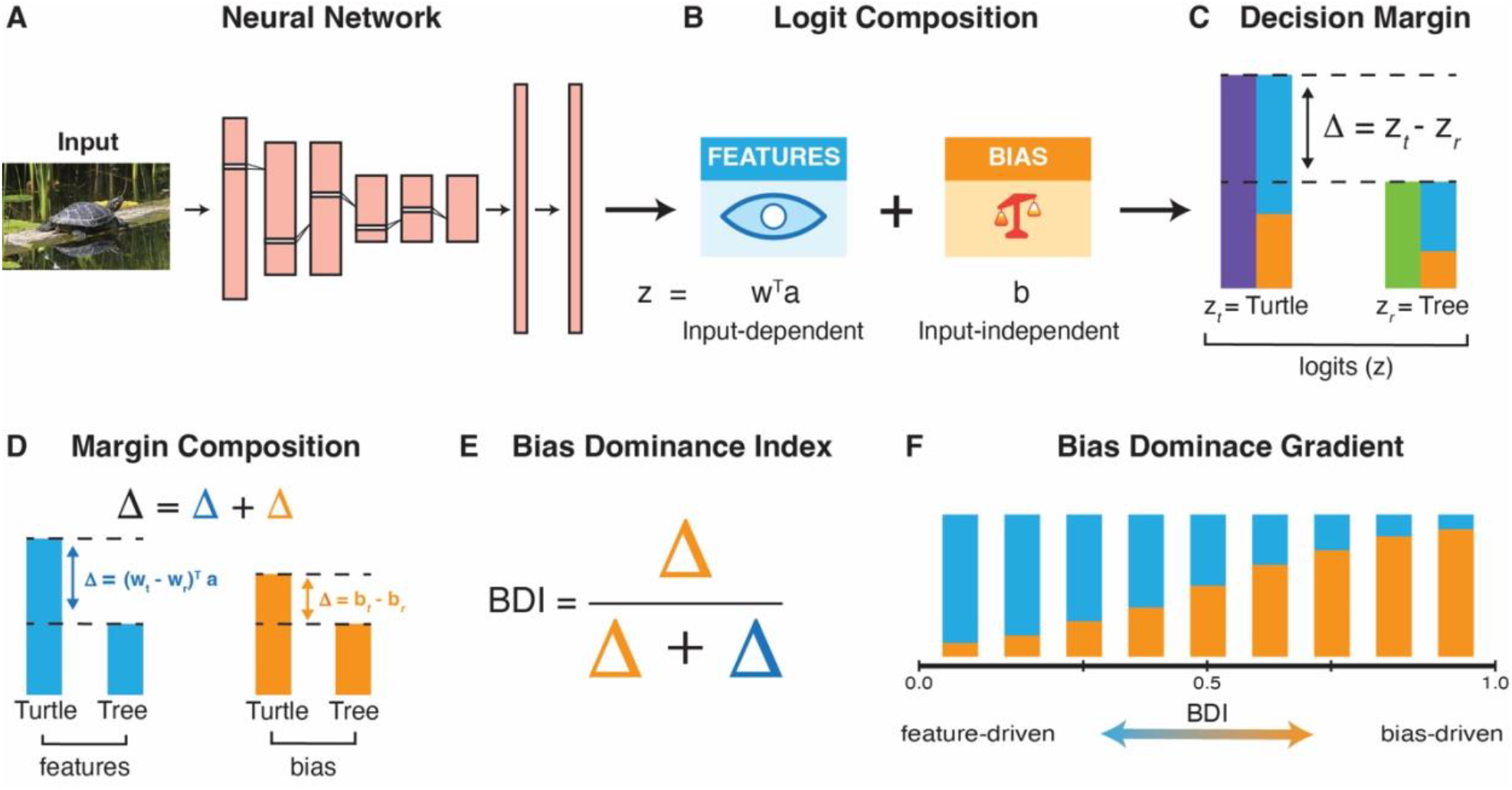
Decision-margin decomposition and bias dominance index. (A) Schematic of a feed-forward neural network that maps an input to an output through successive processing layers. (B) Conceptual separation of the final logit signal into input-dependent feature and input-independent bias components. (C) The decision margin is defined by the logit difference (Δ) between the top predicted class *t* (purple) and its closest competitor *r* (green); blue and orange bars denote the feature and bias contributions to the logits for *t* and *r*, respectively. (D) The decision margin is partitioned into feature and bias contributions for the predicted class *t* and closest competitor *r*. (E) The Bias Dominance Index (BDI) quantifies how much of the margin is supported by bias versus features. (F) Illustration of how BDI varies across inputs/outputs. Low BDI corresponds to feature-grounded margins, whereas high BDI indicates margins that are increasingly bias-supported.

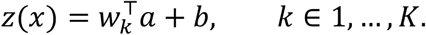

For a target class *t* and alternative class *r*, we define the decision margin as (**Fig. 1C**):

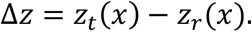

Because the readout is affine, the margin admits an exact decomposition into an input-dependent feature component and an input-independent bias component (**Fig. 1D**):

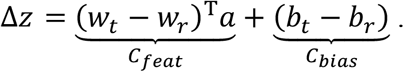

This decomposition isolates not whether the model is confident, but what computationally supports that confidence. Here, “bias” refers specifically to input-independent offsets internal to the model, such as additive terms in affine and normalization operations, rather than to dataset bias or spurious input features.

We use this decomposition to define the Bias Dominance Index (BDI; **Fig. 1E**):

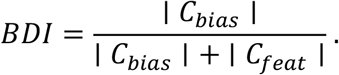

By construction, BDI values near 0 indicate a feature-dominated margin, whereas values near 1 indicate a bias-dominated margin. A value of BDI > 0.5 provides a natural operational criterion for bias dominance, meaning that the input-independent component contributes more to the decision margin than the input-dependent component. Thus, BDI is not a calibration metric or a predictor of accuracy; rather, it provides a mechanism-aware quantification of what supports model decisiveness. Accordingly, BDI quantifies the extent to which the decision margin is supported by these input-independent model- and architecture-intrinsic offsets relative to input-dependent feature-driven computations (**Fig. 1F**).

To map bias-dominant support within the network, we extend this decomposition across depth. For each affine transformation at layer *ℓ*, the pre-nonlinearity output is:

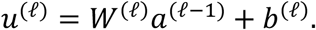

We partition this into an input-dependent feature component:

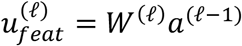

and an input-independent bias component:

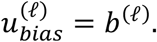

We then compute the gradient of the decision margin with respect to the layer’s pre-nonlinearity output,

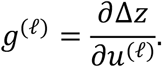

The gradient-weighted feature and bias contributions at layer *ℓ* are:

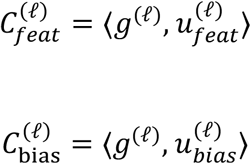

yielding a layer-resolved BDI:

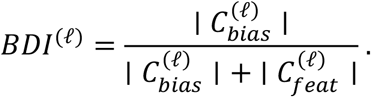

The layer-resolved formulation quantifies the relative alignment of each layer’s feature and input-independent component with the local decision-margin gradient.

Additionally, we define a summary statistic, BDI_all_, that compresses the layer-wise decomposition into a single measure of overall decision composition. For input *i*, we pool the contributions of the bias and feature components across layers by summing 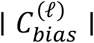 and 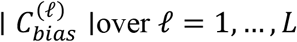:

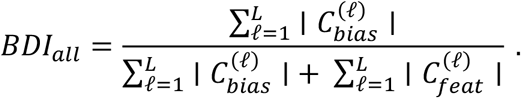

This provides a compact index of the aggregate balance between feature and bias support across the network.

We use standard gradients because BDI is designed as a local decomposition of the current decision margin. The gradient term represents the sensitivity of the margin to each layer output, allowing feature- and bias components to be compared. This provides a computational framework that directly estimates whether the decision margin is supported primarily by input-dependent features or input-independent offsets and is generalizable across model families.

For multiclass settings, we use a label-free version of the framework by defining Δ*z* as the difference between the model’s top-1 and top-2 logits. This allows BDI to be computed without ground-truth labels, making it suitable for auditing deployed systems in which labels are unavailable or delayed. More broadly, the same logic can be extended beyond conventional affine-layer bias terms to other input-independent offsets, such as LayerNorm shifts in vision transformers or additive value-projection terms in transformer language models, enabling a unified analysis of decision composition across architectures (see Methods and Supplemental Information for details). We summarize the input-independent offsets in BDI computation to highlight where bias can enter the computation across model families (**Table 1**).

**Table 1.** Types of input-independent offsets included in the BDI framework.

|  |  |  |  |
| --- | --- | --- | --- |
| Bias Mechanism | Representative Model(s) | Where it enters computation | Rationale for inclusion in BDI |
| Direct additive bias | AlexNet, VGG-16, ViT-b/16, BERT | Convolutional Layers, Dense Linear Layers, Value-projection (attention head) | Standard additive bias term, independent of current input, contributes to the representation and can propagate to the decision margin |
| Normalization offsets | ViT-b/16, BERT | Post-normalization representation (LayerNorm shift, $\beta$) | Standard additive offset term, independent of the current input, that enters the normalized, contributes to the representation and can propagate to the decision margin |

## Results

### BDI reveals decision-composition heterogeneity hidden by confidence

To determine how internal computations support the decision margin, we trained five CNNs on a controlled dot-discrimination task in which each image contained intermixed red and blue dots and the model reported the dominant color (**Fig. 2A**). We previously used this same task in human participants, enabling direct comparison of performance and providing a bridge to human decision-making processes^19^.

**Figure 2.**
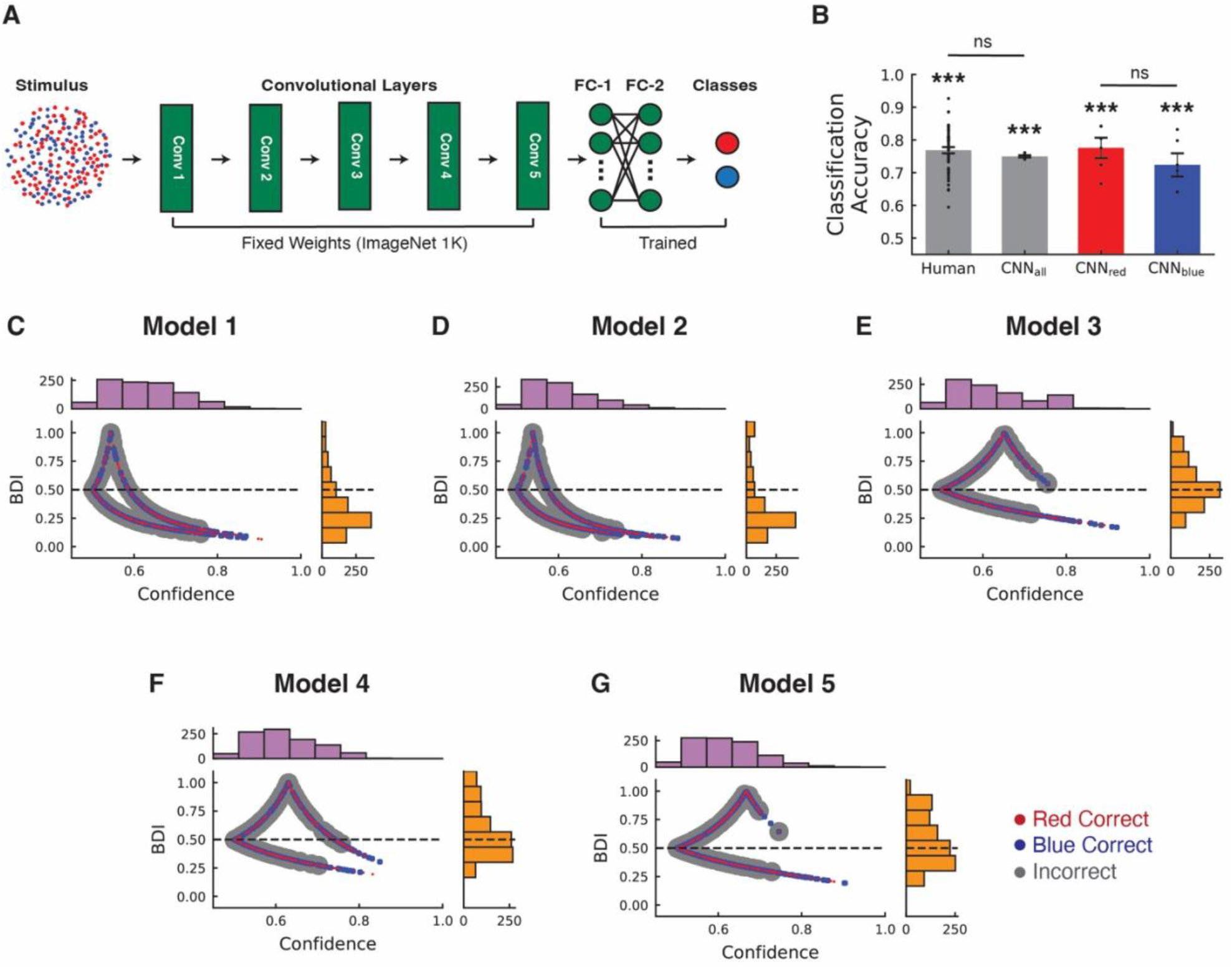
Confidence does not uniquely specify the computational support for the decision in the dot-discrimination task. (A) Schematic of the CNN architecture. During training only the weights of the fully connected layers were modified. Whereas the weights in the convolution component were fixed based on ImageNet. (B) CNN and Human classification performance. Human data is based on 50 subjects from Nakuci et al., 2025 ^19^. Bar plots depict the mean ± sem. Black dots depict the performance of individuals and models, respectively. (C) Relationship between BDI and confidence in Model 1. BDI was computed from the red–blue logit margin at the output affine layer. Confidence is estimated as the *softmax* probability (i.e., the probability assigned to the chosen class) and is neutral at 0.5 for a two-choice perceptual decision task. Each dot corresponds to a trial, colored by outcome. The horizontal dashed line marks the bias-dominance criterion (BDI = 0.5). The histograms summarize the distribution of trials along the confidence and BDI axes. (D-G) Same as panel C, but for Models 2–5, respectively. ***P < 0.001, ns = not significant.

Across models, performance was above chance (Accuracy = 0.75 ± 0.0038; t(4) = 65.41, P = 3.27 × 10^−7^) and comparable to human accuracy on the same task (Accuracy = 0.77 ± 0.0095; t = 1.84, P = 0.07; **Fig. 2B**). Accuracy was above chance for both red-dominant (Accuracy = 0.75 ± 0.035; t(4) = 7.91, P = 1.38 × 10^−3^) and blue-dominant trials (Accuracy = 0.72 ± 0.034; t(4) = 5.65, P = 0.004), with no preference for either color (t(4) = 0.70, P = 0.52).

We then quantified how each model’s decision margin was constructed from the feature component versus the bias component by computing the Bias Dominance Index (BDI) at the last affine layer. Across all five models, the output-layer BDI ranged from weakly to strongly driven by bias (BDI_IQR_ = 0.291 ± 0.036), revealing marked trial-to-trial heterogeneity in how the decision margin was supported (**Fig. 2C-G**). Moreover, on average, 35.48 ± 8.71% of red-dominant trials and 47.80 ± 14.05% of blue-dominant trials exceeded the bias-dominance threshold (BDI > 0.5), indicating that the bias component frequently contributed more to the margin than the feature component. Notably, even for similar *softmax* confidence levels, trials exhibited different BDI values, with some high-confidence responses remaining bias-dominant, demonstrating that decisiveness can coexist with a small contribution from the feature component to the decision margin. These results show that similar confidence values can correspond to different forms of decision support.

### Layer-resolved BDI maps where bias-supported decision components are expressed

Contribution of the bias component is not confined to the output layer; its influence can propagate through the processing hierarchy and shape the decision margin. To quantify how bias dominance evolves with depth, we computed layer-resolved BDI across all convolutional and fully connected layers in the dot-discrimination task, yielding a trial-resolved depth profile of bias dominance. This analysis treats each layer as a potential locus where the bias component can modulate intermediate representations, enabling us to determine how bias- and feature components shape the decision margin.

Across models, layer-resolved BDI showed structured depth dependence (**Fig. 3A–E**). The fraction of bias-dominant trials (BDI > 0.5) was highest in early convolutional layers (BDI_Feature-0_ = 47.10 ± 1.82%; t(4) = 24.87, P_FDR_ = 1.05 × 10^−4^) and re-emerged in late linear layers (BDI_Linear-4_ = 33.14 ± 8.20%; t(4) = 4.04, P_FDR_ = 0.04; BDI_Linear-6_ = 41.66 ± 8.15%; t(4) = 5.11, P_FDR_ = 0.02), indicating that bias-related support can arise upstream and persist to the decision margin (**Fig. 3F**). These profiles identify where offset-supported contributions are expressed.

**Figure 3.**
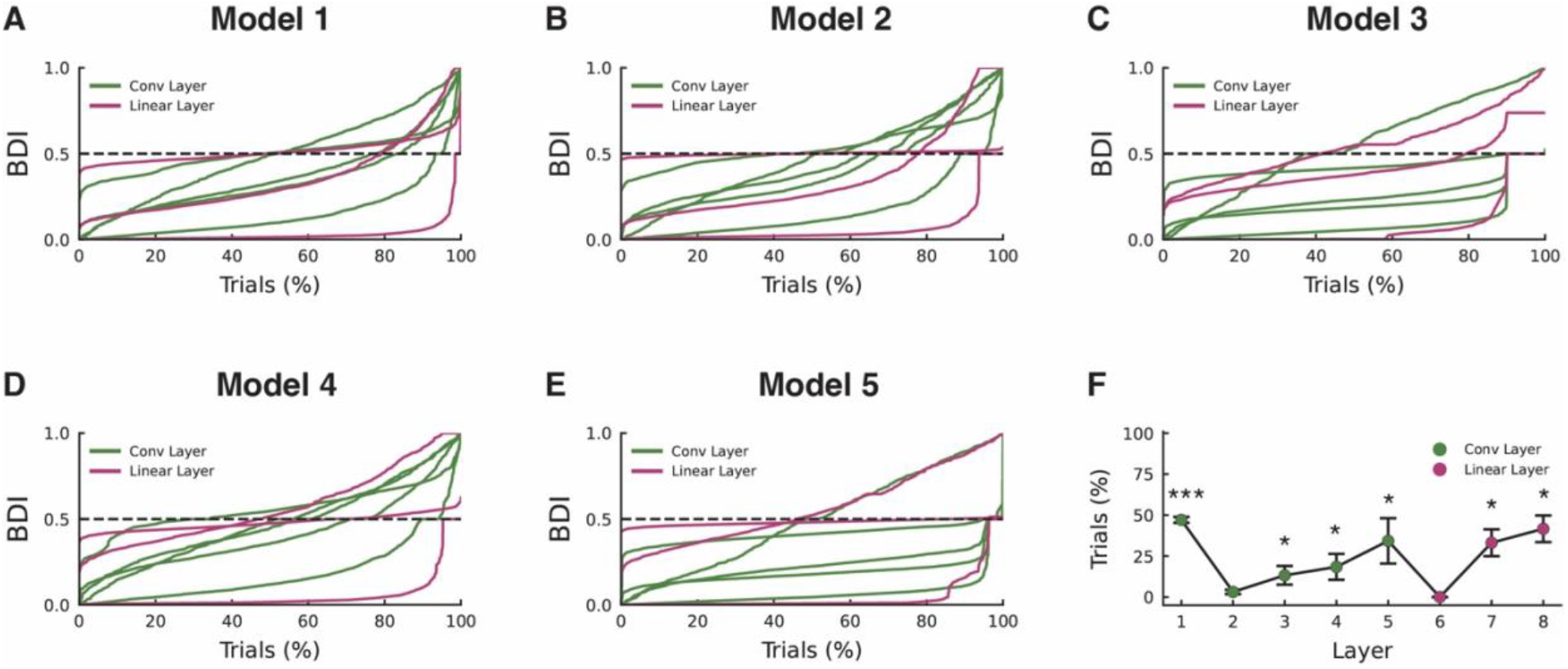
Layer-resolved Bias Dominance Index across network depth in the dot-discrimination task. (A) Layer-resolved BDI profile for Model 1. For each layer, trials are ranked by BDI values. Colored lines correspond to layers spanning convolutional and fully connected linear layers that contain a bias component. The dashed line indicates the bias-dominance criterion BDI = 0.5. (B–E) Same as panel A for Models 2-5. (F) Percentage of trials with BDI > 0.5 across layers. Plot shows Mean ± SEM across models. Layers are ordered sequentially by execution, and only layers with bias terms are included. Statistical significance was estimated using one-sample *t*-test and FDR-corrected. ***P_FDR_ < 0.001 and *P_FDR_ < 0.05.

### Bias support can be stabilizing rather than uniformly harmful

Real-world decision systems must operate under noise and corrupted internal signals. We therefore asked whether the bias component contributes causally to performance when the feature component is degraded. For each test trial, we added Gaussian noise, *N*(0, *σ*), either to the neural activation vector or to the classifier weights mapping the representations to output logits, and quantified accuracy with and without the bias component (*b*_*red*_ = *b*_*blue*_ = 0). Using both perturbations distinguishes sensitivity to representational noise (activation corruption) from sensitivity to readout uncertainty (weight corruption).

In a representative model (Model 4), weight noise impaired accuracy more than activation noise (P_FDR_ < 0.001; **Fig. 4**). To confirm this observation, we tested whether the stabilization from them bias component depended on perturbation magnitude using a three-way ANOVA with noise type (activation vs weights), noise level, and bias condition (with vs without) as factors. We observed significant noise type × noise level – F(9,39960) = 75.95, P = 9.79 × 10^−148^ – and noise type × bias condition interactions – F(1,39960) = 444.20, P = 4.52 × 10^−98^ – indicating that activation and readout-weight perturbations had distinct degradation profiles and were differentially affected by bias. However, the three-way interaction was not significant, F(9,39960) = 0.26, P = 0.98, suggesting that the bias-related stabilization was consistent across the tested noise levels.

**Figure 4.**
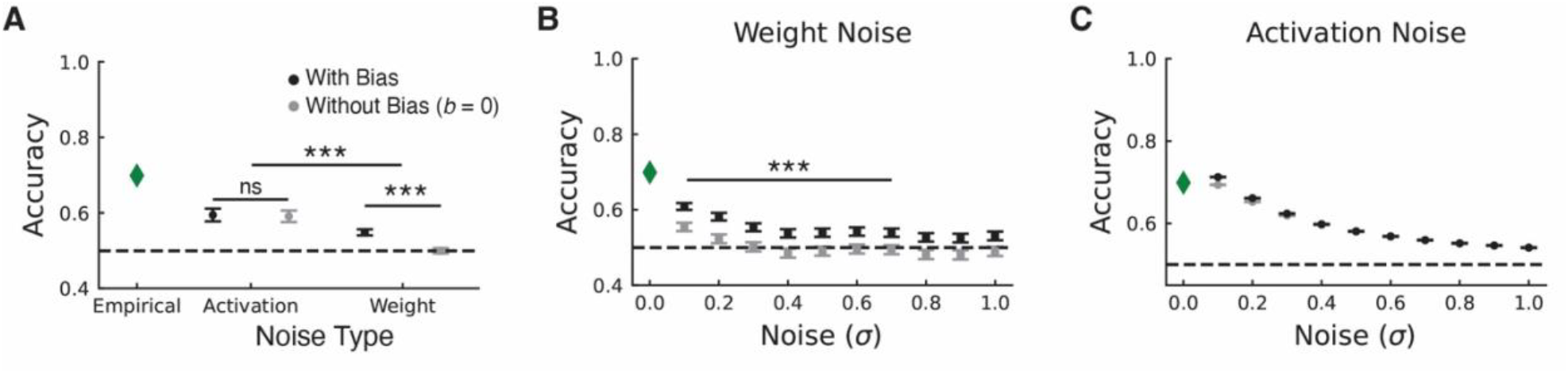
Bias component preserves performance against noise in the weights in the dot-discrimination task. (A) Average classification accuracy after adding noise. Green diamond represents the empirical performance with no added noise (σ = 0) is shown for reference. Black and gray dots represent the average classification value across all noise levels for with and without the bias term (*b*_*red*_ = *b*_*blue*_ = 0), respectively. The dashed line indicates chance performance (0.5). Error bars show Mean ± SEM. (B) Classification accuracy for each level of noise added to the weights. (C) Same as panel B, but noise added to the neuronal activations. Statistical significance was determined using paired-sample *t*-test and FDR corrected. ***P_FDR_ < 0.001; ns = not significant.

Specifically, when noise was added to the weights, retaining the bias component improved performance relative to removing it (Accuracy = 0.55 ± 0.008 vs 0.49 ± 0.006; t(9) = 28.56, P_FDR_ = 3.84 × 10^−10^). Under activation noise, accuracy did not differ with versus without the bias component (Accuracy = 0.59 ± 0.017 vs 0.59 ± 0.015; t(9) = 1.94, P_FDR_ = 0.24). Similar results were observed in the other models, indicating that the bias component can stabilize decisions when readout weights are unreliable (**Fig. S1**). These findings support interpreting BDI as a decision-composition diagnostic rather than a simple error score, because bias support can be beneficial under some perturbations even though its dominance changes the basis on which trust should be assigned.

### BDI provides a label-free axis for auditing pretrained vision models

To assess whether bias dominance extends beyond the controlled perceptual decision-making task, we evaluated ImageNet-pretrained AlexNet, VGG-16, and ViT-b/16 on Tiny ImageNet and Tiny ImageNet-C images. These models produce 1000-way ImageNet logits. We therefore used this setting to determine whether BDI can be used in a label-free manner. Specifically, we quantified the decision support using a label-free margin defined as the difference between the highest and second-highest logits, Δ*z* = *z*_(1)_ − *z*_(2)_, which does not require class labels. Competitor indices (1), (2) were determined per input by the model’s own ranking. We first focused on the relationship between BDI and confidence in the output layer because, in this layer the relationship between the feature component, and the bias component directly influences the decision margin.

Ground-truth labels were not used to define Δ*z* or compute BDI, but labels were used only for post-hoc visualization and mismatch-rate analysis. Label match / mismatch on Tiny ImageNet were computed only after mapping Tiny ImageNet classes (wnid) into the ImageNet-1K (synset) space. Specifically, each Tiny ImageNet wnid is mapped to its corresponding ImageNet-1K class.

In AlexNet, BDI computed from the top-1 versus top-2 logit margin at the affine output layer varied widely across inputs (Mean = 0.114; Std = 0.161; IQR = 0.111; **Fig. S2**). In total, 4.47% of trials exceeded the bias-dominance threshold (BDI > 0.50; **Fig. 5A**). To summarize the relationship between decision composition and *softmax* confidence, we overlaid confidence-binned medians and interquartile ranges of BDI on the trial-level scatter, which revealed a strong inverse trend but substantial within-bin variability at low-to-moderate confidence.

**Figure 5.**
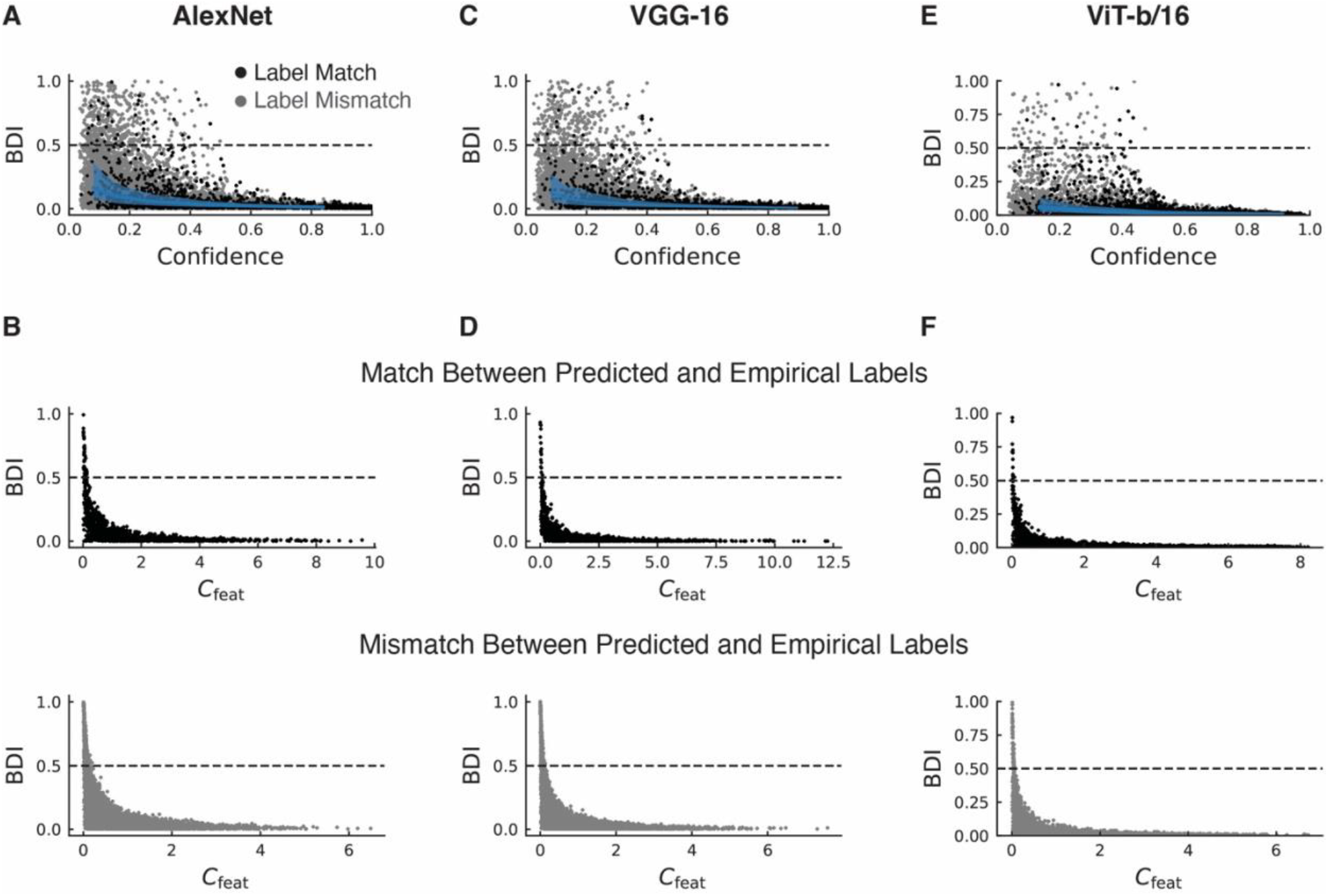
Bias-dominant decision margins persist at high confidence in CNN and ViT models. (A) Relationship between BDI and confidence for AlexNet. BDI was computed from the top-1 versus top-2 logit margin at the affine output layer. Solid blue line shows the median BDI within confidence bins (N_bins_ = 10) and the shaded band indicates the interquartile range, summarizing the central trend and within-confidence variability in decision composition. For visualization, points are colored by label match (black) and mismatch (gray). Confidence is the *softmax* probability assigned to the chosen class (chance baseline 1/1000 = 0.001 for 1000-way ImageNet outputs). Analysis is based on 10,000 images from the Tiny ImageNet dataset. Label match/mismatch is synset-mapped; Tiny ImageNet wnid → ImageNet-1K; used for reporting only. The horizontal dashed line marks the bias-dominance criterion BDI = 0.5. (B) Relationship between BDI and the feature component (*C*_*feat*_). (C,D) Same as panels A and B, but for VGG-16. (E,F) Same as panels A and B, but for ViT-b/16.

Importantly, the confidence–BDI relationship alone does not exclude cases in which the contribution of the feature component, *C*_*feat*_, to the decision margin is large in absolute terms, yet the bias component is larger still. In such cases, BDI remains high despite substantial feature support. We therefore related BDI to *C*_*feat*_, to test whether high BDI primarily reflects reduced feature support for the decision margin, rather than simply an increase in bias magnitude.

When relating BDI to the feature component showed that BDI increased as feature support decreased (**Fig. 5B**). Even in high-BDI/low-*C*_*feat*_ cases, top-1 predictions could still agree with the remapped label, indicating that label agreement can occur despite weak feature support. VGG-16 and ViT-b/16 showed the same qualitative pattern (**Fig. 5C–F; Fig. S2**).

Layer-resolved analyses under the label-free margin mapped bias-dominant support across across network depth. In AlexNet and VGG-16, bias dominance was most pronounced in convolutional layers (**Fig. 6A–D**), whereas in ViT-b/16 it was strongest in LayerNorm modules (**Fig. 6E and F; Fig. S3**). Moreover, to corroborate these findings, on a layer-wise basis we clamped the bias term (*b* = 0) and evaluated the changes in the decision margin and whether the model’s freely chosen top-1 prediction changed (**Fig. S4-S6**). Additionally, we evaluated the BDI index on a subset of images from the standard ImageNet dataset and observed similar results for AlexNet, VGG-16 and ViT_b/16 (**Fig. S7**). These results indicate that bias dominance increases as feature support weakens, even when the predicted label agrees with the mapped ground-truth label, and bias dominance is a depth-distributed computational property across architectures. Together, these findings positioning BDI as a label-free diagnostic for identifying and mapping decisions that are disproportionately bias-supported rather than feature supported.

**Figure 6.**
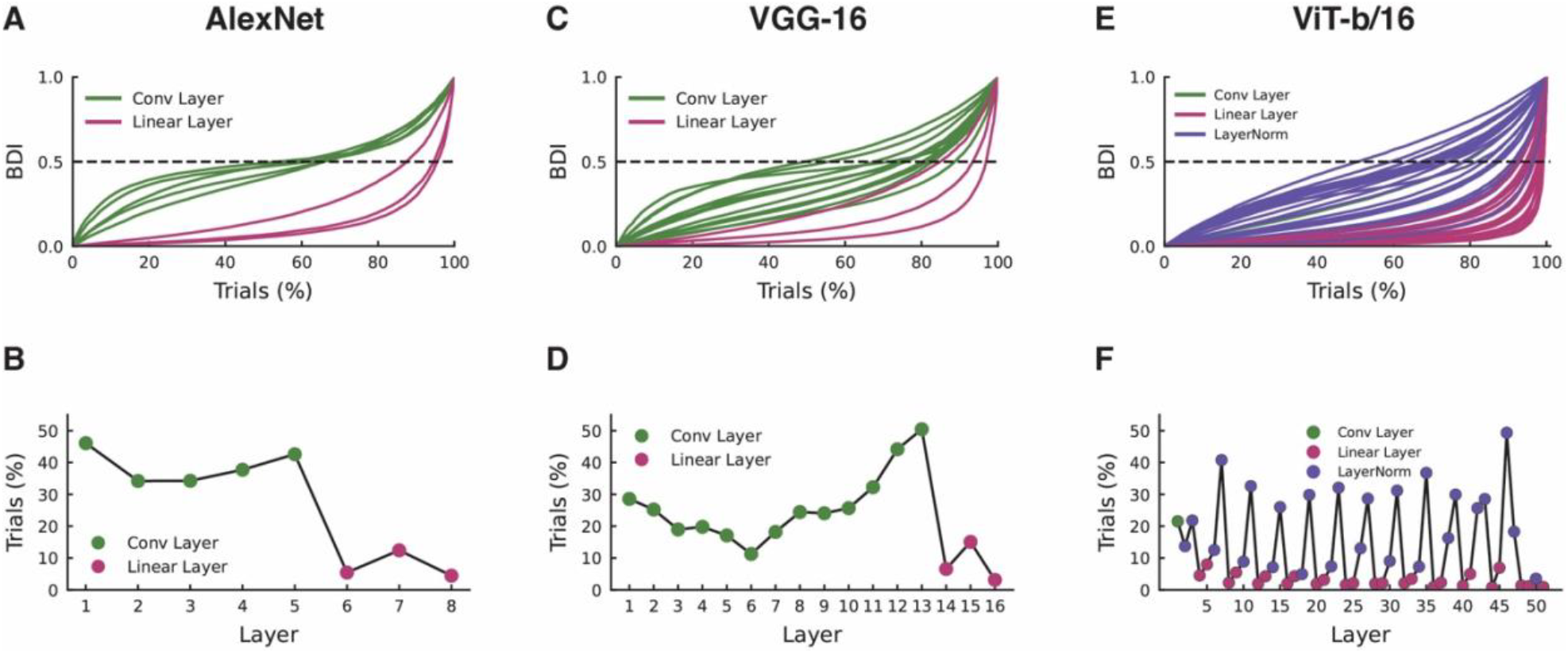
Layer-resolved bias dominance provides a depth-resolved profile of margin support in CNN and ViT models. (A) Layer-resolved BDI profile for AlexNet. Colored lines correspond to layers spanning convolutional and fully connected linear layers that contain a bias component. Analysis is based on 10,000 images from the Tiny ImageNet dataset. Trials are ranked by BDI values. The dashed line indicates the bias-dominance criterion BDI = 0.5. (B) Percentage of bias-dominant trials with BDI > 0.5 across layers for AlexNet. Layers are ordered sequentially according to their execution order on the input, and only layers with bias components are included. (C, D) Same as panels A and B, but for VGG-16. (E, F) Same as panels A and B, but for ViT-b/16.

Having established that BDI captures depth-distributed shifts from feature– to bias–supported margins even in the presence of label agreement, we next tested whether this signature becomes more prevalent under explicit distribution shift induced by corruptions. Under Tiny ImageNet-C corruption, AlexNet again exhibited broad BDI values and an increased prevalence of bias-dominant trials when the logit margin was computed from at the affine output layer (6.80% of trials with BDI > 0.50; **Fig. 7A**). When we matched trials by *softmax* confidence, mismatch rates were comparable across low– and high–BDI strata across confidence bins (t(19) = 0.50; P = 0.62; N = 20 confidence bins; **Fig. 7B**). This does not imply that BDI is redundant; rather, it shows that confidence and accuracy-like outcomes do not identify *how* the model produced a decisive margin. At the same confidence level, the predicted label can be supported primarily by features or by bias, indicating that mismatch-rate and confidence alone are insufficient to determine the dominant decision mechanism on a given trial. The same patterns were observed for VGG-16 (**Fig. 7C and D**) and ViT-b/16 (**Fig. 7E and F**). This dissociation is central to selective trust because two decisions can be matched for confidence and correctness-like outcome while differing in whether the decision margin is primarily feature- or offset-supported.

**Figure 7.**
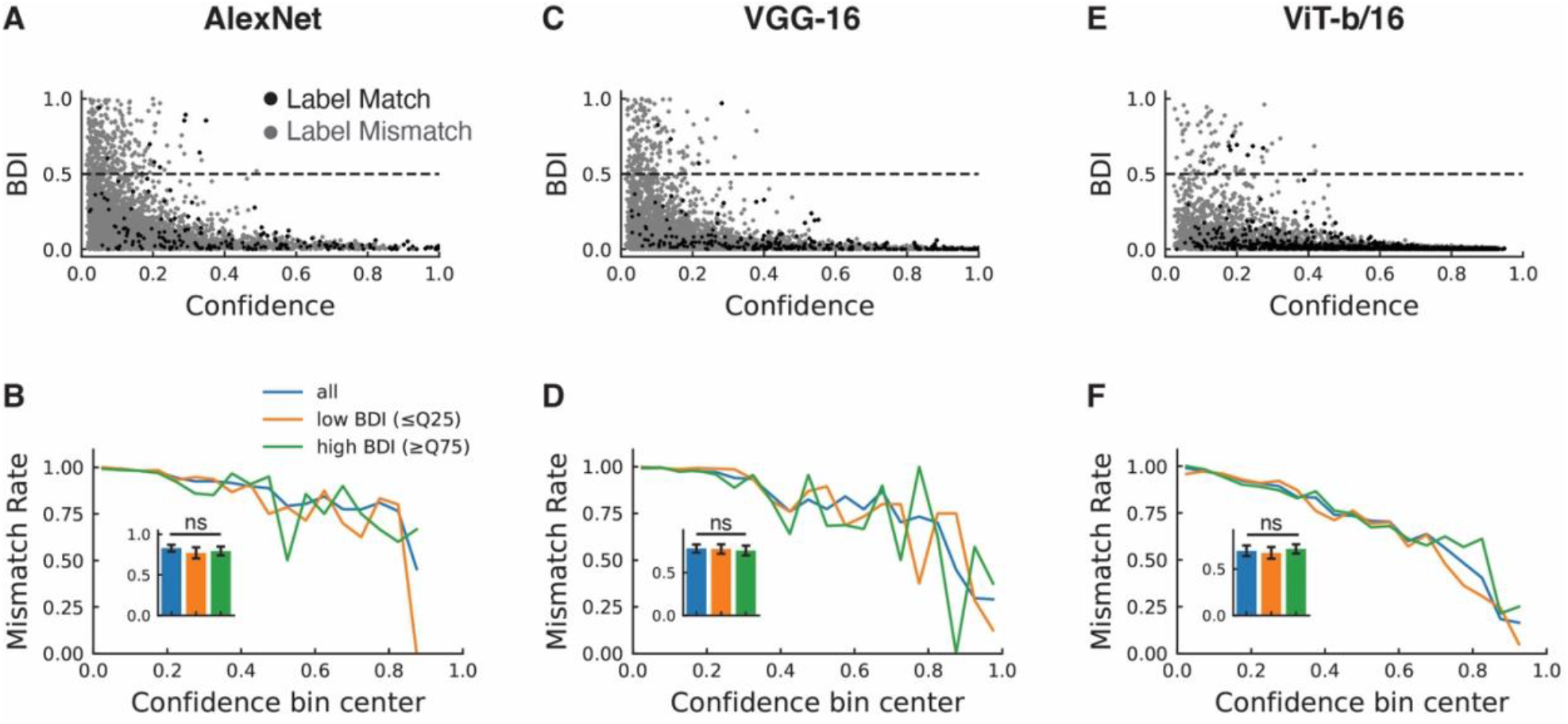
Bias-dominant margins persist under input corruption in CNN and ViT models. (A) Relationship between BDI and confidence for AlexNet on Tiny ImageNet-C. BDI was computed from the top-1 versus top-2 logit margin at the affine output layer. Points are colored by synset-mapped label match (black) versus mismatch (gray) for reporting only (Tiny ImageNet wnid → ImageNet-1K). Confidence is the *softmax* probability assigned to the chosen class (chance baseline 1/1000 = 0.001 for 1000-way ImageNet outputs). Horizontal dashed line denotes the bias-dominance criterion (BDI = 0.5). (B) Label mismatch rate as a function of confidence. Trials are binned by confidence and within each confidence bin trials are partitioned by BDI (bottom vs top quartile), yielding mismatch-rate curves for low-BDI and high-BDI subsets alongside the aggregate (“all”) curve. Inset shows the Mean ± SEM across bins. (C,D) Same as panels A and B, but for VGG-16. (E,F) Same as panels A and B, but for ViT-b/16. Statistical significance was estimated using paired-sample *t*-test. ns = not significant.

### BDI extends decision-composition auditing to language-model token selection

Moreover, to determine whether the BDI framework extends beyond vision models, we applied the analysis to a transformer language model, BERT, in a masked-token completion task^18^. For each prompt, BERT assigns scores to candidate tokens at the masked position, allowing us to define a decision margin as the difference between the highest-scoring token and the next-ranked alternative. Using this margin, we computed layer-resolved BDI across BERT to estimate whether the selected token was supported primarily by input-dependent contextual representations or by input-independent offsets. Across the prompt set, BDI provided a prompt-level measure of decision composition, indicating that the same diagnostic used in CNNs and vision transformers can also be applied to transformer token-selection decisions. Thus, the BERT analysis demonstrates the architectural generality of the framework rather than establishing a new benchmark for language-model reliability.

As can be observed in Table 2, in some prompts, the top-ranked token was contextually plausible, whereas in others it appeared less well matched to the surrounding sentence. Note that that Table 2 provides illustrative examples of this analysis. They show that BDI can be computed for transformer token-selection decisions and inspected alongside the selected token and its competitor. They are not intended as a semantic-coherence benchmark.

**Table 2.** Illustrative examples of BERT completed prompts.

| Table 2. Illustrative examples of BERT completed prompts |  |  |  |
| --- | --- | --- | --- |
| Sample 1 |  |  |  |
| **Prompt:** She reported months of polyuria and polydipsia; HbA1c returned at 11.2%, and the clinician suspected [MASK]. |  |  |  |
| **Completed:** she reported months of polyuria and polydipsia ; hba1c returned at 11. 2 %, and the clinician suspected **infection**. |  |  |  |
| Top-1 Token | Top-2 Token | Top-1 Confidence | BDI Max |
| infection | cancer | 0.0812 | 0.8390 |
| Sample 2 |  |  |  |
| **Prompt:** Although the sky was clear earlier, the sudden gusts and dark clouds suggested an approaching [MASK]. |  |  |  |
| **Completed:** although the sky was clear earlier, the sudden gusts and dark clouds suggested an approaching **storm**. |  |  |  |
| Top-1 Token | Top-2 Token | Top-1 Confidence | BDI Max |
| storm | typhoon | 0.8124 | 0.9844 |
| Sample 3 |  |  |  |
| **Prompt:** The committee agreed that the unexpected results were driven by a hidden [MASK] in the dataset. |  |  |  |
| **Completed:** the committee agreed that the unexpected results were driven by a hidden **flaw** in the dataset. |  |  |  |
| Top-1 Token | Top-2 Token | Top-1 Confidence | BDI Max |
| flaw | vulnerability | 0.2034 | 0.9530 |

Specifically, in prompt 1, the predicted token “infection” appears poorly matched to the surrounding sentence context, which is more strongly aligned with blood glucose levels and diabetes. In contrast, in prompts 2 and 3, the predicted token is contextually appropriate and semantically coherent with the sentence frame. Additional, prompts are provided in Table S1 to confirm that these findings were not limited to these sample prompts. Together, the prompt-level contrast suggest that mechanism-aware analysis may be useful in language models, because the same top-ranked selection process can be bias-driven in both cases, but produce either a context-appropriate completion or a context-mismatched completion.

### Confidence + BDI as a triage policy

To translate decision composition into an operational audit rule, we evaluated a two-parameter, label-free acceptance policy that retains a prediction only when it is both decisive by standard uncertainty criteria and not bias-dominant by the decision-composition diagnostic. This motivates using BDI not only as a descriptive diagnostic, but as a label-free triage axis that targets decision mechanism rather than confidence alone.

Specifically, we accepted trials satisfying Confidence ≥ τ and BDI_all_ ≤ γ, where τ sets a minimum confidence threshold and γ limits tolerance for bias-supported margins. In AlexNet, sweeping (τ, γ) yields the fraction of predictions accepted (**Fig. 8A**). We used this grid to examine the mechanism-aware consequences of acceptance, since lowering γ does not simply reject low-confidence predictions, but selectively excludes decisions whose margins are disproportionately supported by input-independent offsets, including cases that remain high-confidence or match the dataset label. This distinction is important because label agreement alone does not establish that a decision is trustworthy at the mechanistic level. A prediction can be correct while still deriving its decisiveness from weak input-dependent support. Thus, the confidence–BDI policy treats trust as a joint criterion in which accepted decisions should be both confident and feature supported.

**Figure 8.**
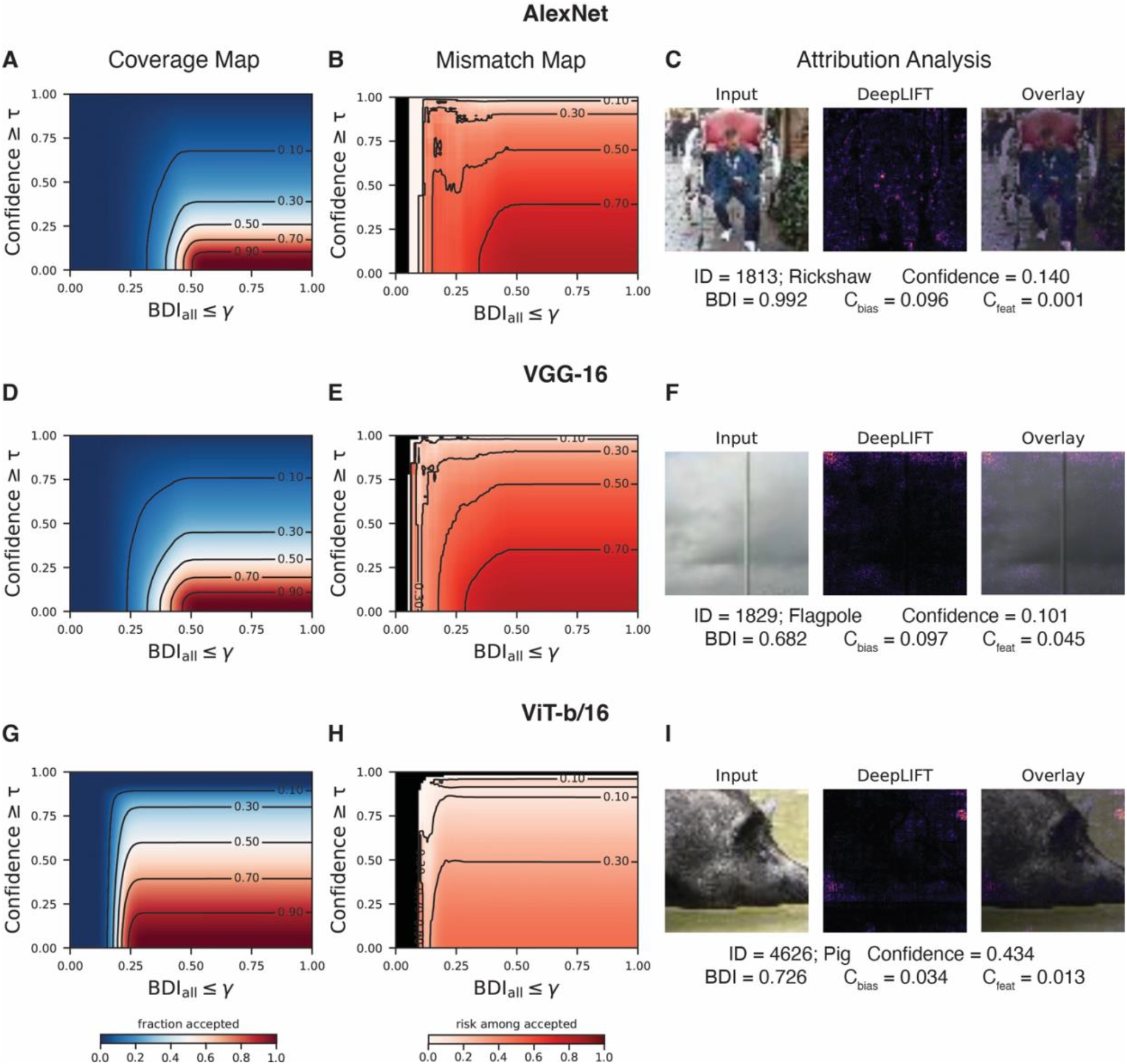
Confidence + BDI acceptance policy for mechanism-aware triage in CNN and ViT models. (A) Coverage heatmaps over the (τ, γ) grid for AlexNet where trials are accepted if Confidence ≥ τ and BDI_all_ ≤ γ. Contours indicate iso-coverage levels. (B) Mismatch-rate among accepted trials. The fraction of accepted predictions that disagreed with the synset-mapped labels. (C) Illustrative trials with high confidence and high BDI value from the output layer whose top-1 predictions mapped to the empirical labels. Despite mapped label match, the decision margin is weakly supported by the feature component (low *C*_*feat*_ relative to *C*_*bias*_), DeepLIFT ^13^ attribution corroborates the limited influence of the feature component on the decision margin. DeepLIFT identifies influential regions, but it remains ambiguous how those regions support the model’s high confidence. Confidence is the *softmax* probability assigned to the chosen class (chance baseline 1/1000 = 0.001 for 1000-way ImageNet outputs). (D-F) Same as panels A-C but for VGG-16. (G-I) Same as panels A-C but for ViT-b/16.

We summarized the outcome of each policy using the mismatch-rate, defined as the fraction of accepted trials whose predicted labels disagreed with the dataset label, producing a corresponding mismatch surface over the same (τ, γ) grid (**Fig. 8B**). At fixed confidence thresholds, stricter BDI thresholds reduced mismatch rates, whereas allowing higher-BDI decisions increased mismatch rates (**Fig. S8**). This provides a decision-level trust interpretation, because confidence indicates that a model is decisive, whereas BDI indicates whether that decisiveness is primarily feature-supported or instead relies disproportionately on input-independent offsets. Confident predictions with low BDI are therefore better candidates for acceptance, whereas confident predictions with high BDI should be treated with caution because their margins are less strongly supported by input-dependent evidence.

Moreover, this policy view fills a critical gap in attribution methods. Attribution methods can highlight regions that influence the output, but they do not directly quantify how the model’s decisiveness is supported, leaving ambiguous how high confidence is warranted when the highlighted features appear weak or misaligned. This is illustrated with a representative high-confidence trial whose predicted label matched the empirical label, yet attribution analysis using DeepLIFT ^13^ did not exhibit a clear correspondence between salient features and either the decision or its confidence (**Fig. 8C**). By contrast, BDI indicated that these decisions were primarily bias-driven, with *C*_*bias*_ dominating *C*_*feat*_. These observations were replicated in both VGG-16 (**Fig. 8D-F**) and ViT-b/16 (**Fig. 8G-I**). Thus, rather than assuming that bias-dominant decisions are always incorrect, the Confidence + BDI policy provides a label-free axis for triage and auditing that can flag confident-but-bias-driven predictions for additional scrutiny.

### BDI measures relative decision support rather than bias magnitude alone

A potential concern is that BDI could be viewed as a re-expression of the bias contribution, since when feature support is attenuated (*C*_feat_ ≈ 0) the index necessarily approaches its bias-dominant limit. However, BDI is not intended to quantify the absolute magnitude of bias, but rather the relative support of the decision margin from bias-versus feature components. To test whether BDI collapses to a bias-only statistic, we computed the bias contribution to the top-1 versus top-2 margin at each layer, 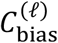, and examined its association with layer-resolved BDI. The relationship between BDI and 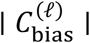 exhibited substantial dispersion in BDI values at fixed 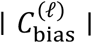 (**Fig. S9–S11**). This pattern indicates that bias dominance is not determined by bias magnitude alone, but depends jointly on bias and feature support, consistent with BDI’s ratio form.

## Discussion

Modern neural networks are often evaluated by whether their predictions are accurate and confident. Here we show that these quantities do not fully determine whether a particular decision should be trusted. Addressing this issue requires moving beyond confidence and the magnitude of the decision margin to examine its composition. This distinction is important for trust because two decisions can have the same confidence values, but their margin could be supported primarily by input-dependent features or by input-independent offsets (i.e. bias). To address this issue, we developed the Bias Dominance Index (BDI), a measure of whether the margin is supported primarily by input-dependent feature contributions or disproportionately by input-independent offsets internal to the model. BDI provides a quantitative lens to see through the “confidence mask” of DNNs. Across large-scale vision and language models – AlexNet^15^, VGG-16^16^, ViT-b/16^17^, BERT^18^ – BDI revealed that confidence can mask mechanistically distinct decisions, showing that predictions with similar apparent confidence may differ in whether their margins are feature-supported or bias-driven.

These findings have practical implications for trusting and auditing modern neural networks. Crucially, accuracy and confidence alone were not a sufficient description of the computational basis of a prediction. Our results indicated that two decisions with similar confidence and similar label agreement can rely on different components. One may be primarily supported by feature-dependent computations, whereas the other may be disproportionately supported by bias. This is consistent with prior work showing that the bias terms can make substantial contributions to prediction accuracy. In certain settings, it has been shown that using only the bias terms can achieve roughly 30–40% of full-model performance^20^.

The influence of bias is not confined to the readout layer but can be mapped to early processing stages and persist through intermediate representations, highlighting that bias is a distributed property of computation. In convolutional neural networks, the influence of bias was strongest in the early layers, indicating that internal priors shape the feature extraction process from its inception.

In biological perception, systematic biases are often interpreted as adaptive priors – shaped by natural stimulus statistics and evolutionary pressures–that help stabilize feature inference under uncertainty^21^. In convolutional neural networks, however, these early offsets may represent architectural shortcuts that allow the model to bypass complex feature synthesis when a prior is sufficiently strong^22,23^. Moreover, in the vision transformer architecture, bias was introduced primarily through layer normalization, pointing to normalization as a key locus where input-independent offsets can be amplified across depth. More broadly, because modern architectures include multiple normalization and other affine operations, the set of loci where input-independent offsets can enter expands, underscoring the need for diagnostic measures. Overall, if one aims to reduce the support of the bias component to the decision margin, effective interventions may need to target early representational dynamics and normalization mechanisms—not only the output-layer decision rule.

While bias in machine learning is often framed as an issue to be corrected^24–26^, our perturbation analysis reveals a more nuanced role. Under conditions of high noise or weight degradation, bias acted as support, preserving classification accuracy when evidentiary weights were compromised. This suggests that the bias in neural architectures may function as a learned internal prior, providing a baseline of robustness in volatile environments. The challenge for future research is to distinguish between *beneficial priors* that aid generalization and *harmful biases* that drive erroneous overconfidence.

Most deployment pipelines treat *softmax* confidence as a proxy for reliability and implement a one-dimensional acceptance rule, for example, accepting predictions only if confidence exceeds a threshold^27^. Our results suggest that confidence alone can conflate qualitatively different factors underlying decisions. Predictions can be equally confident yet differ in whether the decision margin is supported by input-dependent features or by input-independent bias. Instead, a two-parameter policy that jointly thresholds confidence and BDI could allow practitioners to trade coverage against tolerance for bias-dominant margins. Because both quantities are computed from internal decision variables, such a policy is label-free and could be deployed when ground truth labels are unavailable or delayed. More broadly, this perspective reframes “reliability” as not only a question of margin magnitude (confidence) but also of margin composition (feature vs bias), providing an interpretable handle for auditing and triaging predictions that would otherwise pass confidence screening.

BDI can also be estimated with label-free inputs (top-1 vs. top-2) expanding the practical utility of this framework. BDI could be applied in high-stakes deployments (e.g., medical diagnostics^28^) where ground truth is unavailable or delayed. In such settings, BDI could serve as a monitor that flags highly confident decisions whose margins are primarily bias-supported. This may provide a complementary audit signal for identifying predictions that warrant additional review. In this way, BDI could help flag cases at risk of overconfident errors, spurious predictions, over-extrapolation, and brittle generalization.

Beyond machine learning, these findings may also inform computational neuroscience models of the visual cortex and decision-making. Our observation that bias-dominant margin support can emerge early and persist across depth may be a way to formalize how *priors* influence perceptual decisions when stimulus-linked support is weak. This framing may be relevant because CNNs have been widely used as mechanistic models for multiple aspects of human vision, including object recognition^29,30^, representational structure^31^ and processing in the visual cortex^32–34^.

Moreover, in prior work, humans performing the same dot-discrimination task used in the current analysis exhibited multiple brain activation patterns, including engagement of the Default Mode Network^19^. Default Mode Network engagement is associated with brain states in which internally generated dynamics dominate over externally driven dynamics^35^. In this light, our results raise the possibility that these pattern shifts reflect changes in decision composition.

Accordingly, we speculate that Default Mode Network engagement during decision-making may be preferentially associated with trials in which sensory features provide relatively little margin support and internal priors provide comparatively greater support, paralleling bias-driven decision margins in DNNs. While we are not able to test this relationship here, future work could evaluate the parallel between bias-driven decisions in DNNs and Default Mode Network engagement in humans.

Several limitations of the present work should be noted. One limitation of the current BDI formulation is that it captures only the input-independent component, and not all forms of undesirable reliance in a decision. In particular, BDI does not by itself identify other spurious but input-dependent biases, including bias that may be encoded in the learned weights or feature representations rather than in explicit additive offsets. Future extensions of the BDI framework could therefore broaden the analysis from input-independent support to more general forms of non-evidence-grounded support. In addition, the current study focused primarily on ImageNet-trained vision models, and future work should assess BDI on a wider range of state-of-the-art baselines, particularly for large language models, to provide a more detailed evaluation of specific model classes. Finally, the analysis was centered mainly on common corruptions and noise. Therefore, future studies should test BDI across a broader range of domains and under adversarial perturbations to determine whether bias dominance is systematically linked to model robustness.

By shifting the focus from input features to decision composition, we provide a foundation for auditing neural networks. Importantly, this framework can be applied to any model whose decision variables can be expressed as affine functions of internal representations. Moreover, future work tracking the developmental trajectory of BDI over training could pinpoint when a model transitions from learning input-driven structure to exploiting bias. Taken together, rather than merely rationalizing outputs post hoc, the BDI framework exposes the balance between input-linked support and model-intrinsic offsets, moving us toward AI systems whose decisions are not only accurate, but more readily interpretable.

## Methods

### Decomposition of the decision margin and bias dominance index

For each input *x*_*i*_ (trial *i*), logits are produced by an affine readout from an internal activation vector *a*_*i*_ from a total of *K* classes:

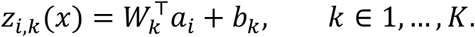

We define the logit margin between a target class *t* and alternative class *r* as:

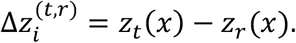

The margin admits an exact decomposition into an input-dependent feature component 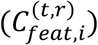 and an input-independent bias component 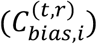:

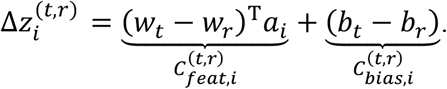

We define the Bias Dominance Index (BDI) for the decision margin between (*t, r*) on trial *i* as:

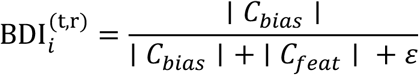

with *ε* > 0 for numerical stability. By construction, BDI ∈ [0,1] values near 0 indicate feature-dominant margin, whereas values near 1 indicate margins dominated by bias. We use BDI > 0.5 as a natural dominance threshold. Unlike margin magnitude or confidence alone, BDI isolates how a margin is constructed.

#### Layer-resolved BDI

To extend BDI beyond the output layer, an analogous feature–bias decomposition at multiple stages of the network was computed. This is accomplished by extending BDI across depth through attributing each layer’s feature and bias contributions to the decision margin. Specifically, for each affine transformation at layer *ℓ*, the pre-nonlinearity output is:

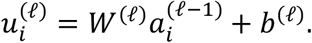

We partition the output into an input-dependent feature component:

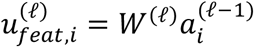

and a bias component (broadcast across spatial positions for convolutional layers):

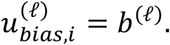

In convolutional neural networks (CNNs), the input-independent component, 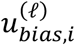 corresponds to the additive bias parameters in affine layers. In the vision transformer network, ViT-b/16, there is an additional input-independent bias component (i.e. offset) in the LayerNorm module, the *β* parameter. We treat the LayerNorm shift (β) as an input-independent offset because it contributes an additive term to the post-normalization representation that does not depend on the current input sample. In the label-free decision margin Δ*z* = *z*_(1)_ − *z*_(2)_, the contribution of *β* can be written as Δ*z*_LN_ = (*w*_(*t*)_ − *w*_(*r*)_)^⊤^*β*. Note that, the Δ*z*_LN_ term is independent of the current input, but is conditional on the trial’s competitor classes (1) and (2). When computing label-free margin support (top-1 versus top-2), offsets are propagated through subsequent affine projections and can contribute to the decision margin even in the absence of strong input-linked support. Accordingly, we include β-derived contributions in the bias component for ViT (derivation in the Supplemental).

In BERT self-attention, the value-projection bias introduces an input-independent contribution to each attention head output. For head *j*, the head output at token *t* is 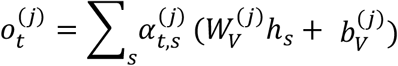. Because the attention weights are normalized 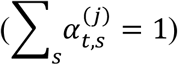, this decomposes as 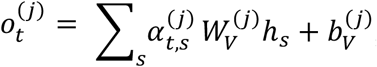, showing that 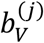 is transmitted as an additive, input-independent offset. After head concatenation and output projection, this yields a block-level additive term 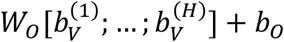, which can contribute directly to the decision margin and therefore to the layer-resolved Bias Dominance Index (BDI). By contrast, query- and key-projection biases (*b*_*Q*_, *b*_*K*_) act primarily through the attention scores and hence modulate the attention weights 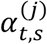 (that is, routing/selection), rather than contributing a simple additive, input-independent offset to the head output representation (derivation in the Supplemental). Note that *b*_*Q*_ and *b*_*K*_ are included here for completeness, but these forms of bias are not used in the BDI analysis.

The gradient of the output margin with respect to the layer’s pre-nonlinearity activations is defined as:

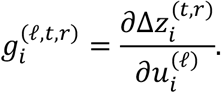

The gradient-weighted feature and bias contributions at layer *ℓ* is estimated as:

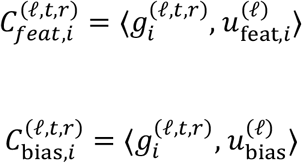

where ⟨⋅,⋅⟩ denotes the inner product (includes all spatial positions for convolutional layers). The layer-resolved BDI is:

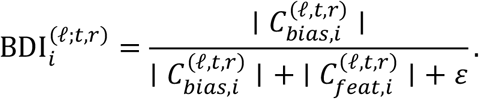

The layer-resolved formulation quantifies the relative alignment of each layer’s input-dependent and input-independent component with the local decision-margin gradient.

Additionally, we define a summary statistic, BDI_all_, that compresses the layer-wise decomposition into a single measure of overall decision composition. For input *i*, we pool the contributions of the bias and feature components across layers by summing 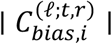 and 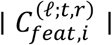 over *ℓ* = 1, …, *L*:

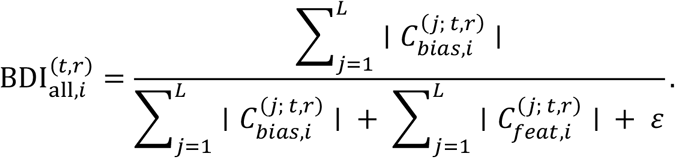

This provides a compact index of the aggregate balance between feature and bias support across the network.

The gradient-based procedure used to estimate layer-resolved BDI may be sensitive to local saturation^36^ and therefore may understate support when derivatives are small ^12^. BDI decomposes the current decision margin into input-dependent and input-independent support terms. By contrast, Integrated Gradients (IG) attributes the change in output from a baseline input to the current input along a path and therefore measures a different quantity. IG is not a direct replacement for BDI because the two methods attribute different quantities. Because IG is defined on baseline-relative differences, explicit input-independent offsets cancel from the attributed output difference and are therefore not directly recoverable as a separate bias-support term. Thus, an IG-based extension would not be a direct replacement for BDI, but rather a distinct path-integrated variant that may provide complementary information. Nevertheless, future analyses could evaluate if IG-weighted BDI provides additional information to aid in diagnosis and triaging.

#### Competitor selection

The most generalizable form of the analysis is to estimate BDI on each input between the primary and secondary choices because this does not require a ground truth and can be applied to cases with multiple classes. Specifically, the primary choice is the predicted class *t* = *argmax*_*k*=*t*_(*z*_*k*_) and the class associated with the second-largest logit *r* = *argmax*_*k*≠*t*_(*z*_*k*_). This defines BDI on the smallest “winning margin” separating the model’s top prediction from its nearest alternative. If the ground truth is known, then the target, *t*, is the logit from the ground-truth class and the alternative, *r*, is the highest-logit from the incorrect class. This yields a label-anchored measure of whether correct classification is supported primarily by the feature or by the bias component. In the special case of two classes, competitor selection is fixed by the task.

### Dot-discrimination task

The convolutional neural network (CNN) was trained to perform a perceptual decision-making task, identifying whether red or blue dots were more frequent in a cloud of randomly distributed dots. Specifically, the CNN was trained to perform a modified version of a perceptual decision-making task we have previously used in humans to understand the neural correlates of decision-making^19^. Each stimulus consisted of either 90 red- or 100 blue-colored dots, presented within an imaginary circular aperture. Dot positions were randomized for every trial to minimize low-level visual regularities. For testing, 1,000 novel stimuli were created with equal numbers of red- and blue-dominant trials. All images were generated using Python.

### Dot-discrimination task CNN model, training and testing

We used a CNN based on the AlexNet^15^ architecture, modified for binary classification. Convolutional-layer weights were fixed to ImageNet-1K–pretrained parameters provided by PyTorch (https://pytorch.org/), and only the fully connected layers were trained. This design keeps the feature-extraction stage stable while allowing the decision stage to adapt to the red/blue task. Using pretrained weights learned from diverse natural images to isolate decision-level effects from feature learning.

The CNN was trained on randomly generated trials of the dot-discrimination task. Training was conducted in five batches of randomly chosen trials. Model optimization was performed using stochastic gradient descent as implemented in PyTorch. During training and testing, images were balanced for red and blue dominance. After training, the CNN was evaluated on 1,000 unique trials not included in the training set. Accuracy was assessed separately for red-dominant and blue-dominant images, as well as across all trials. In total five different CNN models were trained and tested. Training and testing were conducted in Google Colab on an NVIDIA A100 GPU.

### Dot-discrimination task noise perturbations of input representations and readout

In the dot-discrimination task, for each trial in the test set, noise was applied either to the activation pattern or to the readout weights, thereby perturbing the input representation or its mapping to the output, respectively. Activation noise was modeled by adding Gaussian noise, *N*(0, *σ*), to the neuronal activations in the penultimate layer, which reflect the network’s internal representation of the input. Weight noise was modeled by adding Gaussian noise, *N*(0, *σ*), to the weights that mapped activations to the classification output. The standard deviation of the noise (*σ*) ranged from 0.1 to 1.0, and classification accuracy was estimated from 1,000 simulations at each noise level with and without the bias terms.

### Tiny ImageNet evaluation with ImageNet-pretrained networks

To assess the generality of the analysis beyond the dot-discrimination task, we evaluated pretrained convolutional neural networks on images from the ImageNet dataset (Tiny ImageNet-200, a 200-class subset of ImageNet^37^). We analyzed all 10,000 images from the held-out validation set. Images were resized and center-cropped to 224×224 and normalized using the standard ImageNet-1K preprocessing associated with the pretrained model weights. Network outputs were the 1000-way ImageNet-1K logits. When we report “label match/correctness,” it denotes whether the model’s top-1 predicted index equals the Tiny ImageNet class index for that image; this serves only as a post-hoc reliability annotation and is not interpreted as semantically aligned ImageNet accuracy. Label match/mismatch refers to synset-mapped agreement between the model’s top-1 ImageNet-1K prediction and the mapped ImageNet-1K label for Tiny ImageNet.

### Layer-wise bias-clamp analysis

To test whether layer-resolved BDI corresponded to finite effects on the final decision, we performed a layer-wise bias-clamp perturbation analysis. For each eligible bias-containing layer, including convolutional, linear and LayerNorm offset terms, we scaled that layer’s bias parameter to zero while leaving all other model parameters unchanged. We then recomputed the model output for the same images and evaluated the original baseline top-1 versus top-2 margin, keeping the baseline top-1 and top-2 classes fixed. The primary perturbation metric was the change in this fixed margin:

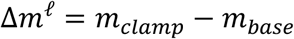

where *m*_*base*_ is the original margin and *m*_*clamp*_ is the margin after clamping the bias of layer *ℓ*. Where Δ*m <* 0 indicates that clamping reduced the original top-1 versus top-2 margin, Δ*m* > 0 clamping increased the margin and Δ*m* = 0 indicates no change. We also quantified whether the model’s freely chosen top-1 prediction changed.

### ImageNet-C evaluation under corrupted inputs

To test whether decision composition persists when feature input is degraded, we evaluated ImageNet-pretrained networks on corrupted inputs from Tiny ImageNet-C, a corruption benchmark constructed by applying ImageNet-C-style perturbations to Tiny ImageNet images^38^. We focused on Gaussian noise corruptions at five severity levels (1–5). AlexNet^15^, VGG-16^16^, and ViT-b/16^17^ were evaluated in inference mode using the canonical ImageNet preprocessing associated with each model’s pretrained weights (resize/crop and per-channel normalization). The analysis was based on a random sample of 5,000 Tiny ImageNet images corrupted with Gaussian noise, with 1,000 images drawn at each severity level.

For each corrupted input, we defined a label-free decision margin as the difference between the highest and second-highest logits, Δ*z* = *z*_(1)_ − *z*_(2)_, where (1) and (2) index the model’s top-1 and top-2 predicted classes for that image. Confidence was quantified as the *softmax* probability from the top-1, *confidence* = *softmax*(*z*_1_), corresponding to the probability assigned to the chosen class (neutral at 1/1000 = 0.001 for a 1000-way decision). Bias Dominance Index (BDI) at the output layer was computed in the same manner from the input-dependent feature component and an input-independent bias component.

To test whether decision composition distinguishes mechanisms beyond what is identifiable from confidence alone, we matched trials by confidence and then stratified by BDI. We stratified images into confidence bins spanning [0,1] (bin_width_ = 0.05; for a total of 20 bins) and, within each bin, split trials into low– and high–BDI subsets using a quantile criterion (bottom vs top quartile). This matched-confidence design controls for confidence by construction and isolates variation attributable to decision composition. When reporting mismatch rates, correctness (synset-mapped) was defined as whether the model’s top-1 ImageNet-1K prediction matched the ImageNet-1K class index obtained by mapping the image’s Tiny ImageNet WordnetID into the ImageNet-1K synset list. This mapping is used only for reporting; BDI and confidence are computed without labels.

### Confidence-BDI acceptance gating and mismatch-rate

We operationalized decision composition as a two-parameter acceptance gating policy that retains a prediction only when it is both decisive by standard uncertainty criteria and feature-grounded by the decision-composition diagnostic. For each trial *i*, we computed confidence, \Conf_*i*_, as the *softmax* probability assigned to the model’s top-1 prediction, and computed BDI_all,*i*_ as the aggregate bias-dominance statistic. Given thresholds (τ, γ), we defined the accepted set as:

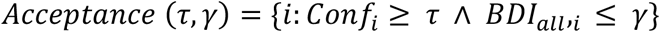

where τ sets a minimum confidence requirement and γ limits tolerance for bias-supported margins. Sweeping (τ, γ) over a uniform grid produced an acceptance surface whose value at each point is coverage, Coverage(*τ, γ*) = ∣ *Acceptance*(*τ, γ*) ∣/*N*, the fraction of predictions accepted (visualized as heatmaps with iso-coverage contours).

To quantify the mismatch-rate associated with each policy, we defined a per-trial indicator *R*_*i*_ ∈ {0,1} that equals 1 for a mismatch and 0 otherwise, and computed the mismatch-rate score (MRS) as the conditional mean over accepted predictions as:

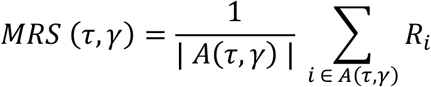

with policies that accept no predictions assigned undefined mismatch-rate (NaN). This policy should not be interpreted as a conventional uncertainty baseline whose sole purpose is to minimize mismatch rate. Rather, it defines a mechanism-aware triage rule in which, among predictions that are sufficiently confident, γ controls tolerance for margins that are disproportionately supported by input-independent offsets.

## Data availability

Trained CNN models are available at https://osf.io/efnq6/. Tiny-ImageNet data can be downloaded at http://cs231n.stanford.edu/tiny-imagenet-200.zip.

## Code availability

The analysis was based on a combination of publicly available toolboxes and analysis specific scripts. Specifically, the CNNs used in the perceptual decision-making task were based on the AlexNet architecture from the PyTorch library (https://pytorch.org/). Trained AlexNet, VGG-16, and ViT-b/16 can be downloaded from the PyTorch library (https://pytorch.org/). Trained CNN models and scripts for formal analysis are available at https://osf.io/efnq6/. A lightweight `ComputeBDI` function for computing aggregate and layer-resolved Bias Dominance Index estimates for individual model decisions is available at https://github.com/jnakuci/Bias-Dominance-Index.

## Declaration of generative AI and AI-assisted technologies in the writing process

During the preparation of this work the author used ChatGPT-5.2 in order to improve the readability and language of the manuscript. After using this tool, the author reviewed and edited the content as needed and takes full responsibility for the content of the published article.

## Acknowledgments

We would like to thank Dobromir Rahnev and Marshall Green for their suggestions. This research was supported by the U.S. Army DEVCOM Army Research Laboratory through mission funding (JOG) and army educational outreach program (JN, W911SR-15-2-0001). The views and conclusions contained in this document are those of the authors and should not be interpreted as representing the official policies, either expressed or implied, of the U.S. Army DEVCOM Army Research Laboratory or the U.S. Government. The U.S. Government is authorized to reproduce and distribute reprints for Government purposes notwithstanding any copyright notation herein.

## Author Contributions

Conceptualization: JN

Methodology: JN

Data Curation: JN

Visualization: JN

Funding Acquisition: JN

Writing – original draft: JN

Writing – review & editing: JN

