## Supplemental Information for "A layer-resolved diagnostic identifies bias-driven decisions in deep neural networks"

Johan Nakuci[1*]

[1]U.S. Army DEVCOM Army Research Laboratory, Aberdeen Proving Ground, Aberdeen, MD, 21005, USA

### Supplemental Methods

Stimulus independent offsets from LayerNorm in transformer models

In transformer architectures, stimulus independent contributions to the decision margin can arise not only from explicit affine biases, but also from normalization. For a hidden representation $h \in \mathbb{R}^d$, LayerNorm is defined as

$$\mathrm{LN}(h) = \gamma \odot \frac{h - \mu(h)}{\sigma(h)} + \beta$$

where $\mu(h)$and $\sigma(h)$ are the mean and standard deviation computed across the $d$ hidden units of a token, $\gamma$ is a learned scale, and $\beta$ is a learned shift. Critically, $\beta$ is a fixed parameter that does not vary with the current input and therefore constitutes a stimulus independent offset at the output of the normalization module.

To see how this offset can propagate into the decision margin, consider a downstream affine map $y = W\,\mathrm{LN}(h) + b$. Substituting the LayerNorm definition yields:

$$y = W\left(\gamma \odot \frac{h - \mu}{\sigma}\right) + (W\beta + b)$$

so that $W\beta$ acts as an effective bias term at the output of the module. At the output of the LayerNorm module, this separates into:

$$u_{\text{feature}} = \gamma \odot \frac{h - \mu(h)}{\sigma(h)}$$

and

$$u_{\text{bias}} = \beta.$$

For the label-free margin used throughout the main text, $\Delta z = z_{(1)} - z_{(2)}$, the LayerNorm shift contributes:

$$\Delta z_{\text{LN-bias}} = (w_{(1)} - w_{(2)})^{\top}\beta$$

where $w_{(1)}$and $w_{(2)}$ are the readout rows for the top-1 and top-2 logits on that trial. This contribution is stimulus-independent conditional on the selected $(1),(2)$ and is therefore grouped with other stimulus-independent terms in our decomposition.

**Accounting for LayerNorm in layer resolved BDI.** In ViT-b/16, we quantify stimulus independent contributions from (i) affine bias parameters in linear projections and the classifier head and (ii) LayerNorm shift parameters $\beta$. Operationally, we treat each LayerNorm module as contributing a bias term given by its $\beta$, and propagate this contribution to the final decision margin using the same layer resolved attribution pipeline used for affine layers. In contrast, the LayerNorm scale parameter $\gamma$ is multiplicative and is not classified as a bias term in our framework.

The set of modules included in the ViT-b/16 layer resolved analysis comprised the initial patch embedding convolution, linear layers within attention and MLP blocks that contain bias parameters, the classifier head, and all LayerNorm modules.

Query/key/value decomposition of attention-head contributions to the decision margin

Consider one self-attention head $j$ in a BERT layer. For clarity, we omit the head index $j$ until the multi-head aggregation step. Let $h_s \in \mathbb{R}^{d_{\text{model}}}$denote the input representation at source token $s$, and let $t$ denote the query token. Define affine query, key, and value projections:

$$q_t = W_Q h_t + b_Q,$$

$$k_s = W_K h_s + b_K,$$

$$v_s = W_V h_s + b_V.$$

The attention score and attention weights are:

$$a_{t,s} = \frac{q_t^\top k_s}{\sqrt{d_h}},$$

$$\alpha_{t,s} = \mathrm{softmax}_s(a_{t,s}),$$

where $d_h$is the head dimension. The head output is:

$$o_t = \sum_s \alpha_{t,s}\, v_s.$$

Substituting the affine forms for $q_t$and $k_s$:

$$q_t^\top k_s = (W_Q h_t + b_Q)^\top (W_K h_s + b_K).$$

This expands to:

$$q_t^\top k_s = h_t^\top W_Q^\top W_K h_s \ + \ b_Q^\top W_K h_s \ + \ h_t^\top W_Q^\top b_K \ + \ b_Q^\top b_K,$$

and substituting in the expanded form yields:

$$a_{t,s} = softmax\left(\frac{h_t^\top W_Q^\top W_K h_s}{\sqrt{d_h}} + \frac{b_Q^\top W_K h_s}{\sqrt{d_h}} + \frac{h_t^\top W_Q^\top b_K}{\sqrt{d_h}} + \frac{b_Q^\top b_K}{\sqrt{d_h}}\right).$$

For fixed query token $t$, $h_t^\top W_Q^\top b_K$ and $b_Q^\top b_K$ are constant and cancel out within *softmax.* On the other hand, $b_Q^\top W_K h_s$ varies with $s$, so it can change attention weights, but it is still input-dependent through $h_s$. Therefore the $a_{t,s}$ reduces to:

$$a_{t,s} = softmax\left(\frac{h_t^\top W_Q^\top W_K h_s}{\sqrt{d_h}} + \frac{b_Q^\top W_K h_s}{\sqrt{d_h}}\right).$$

So $b_Q$ can encode a *prior-like preference* over keys via $b_Q^\top W_K h_s$, but it is not a clean "additive offset" term at the representation level.

**Value bias produces a direct additive offset in the head output.** Expanding the head output using $v_s = W_V h_s + b_V$:

$$o_t = \sum_s \alpha_{t,s} (W_V h_s + b_V) = \sum_s \alpha_{t,s} W_V h_s + \sum_s \alpha_{t,s} b_V.$$

Because $\sum_s \alpha_{t,s} = 1$ as defined by *softmax*, then $o_t$ reduces to:

$$o_t = \sum_s \alpha_{t,s} W_V h_s + b_v.$$

This shows that $b_V$ is transmitted as a direct additive, input-independent offset in the head output. Whereas the query bias contribution ($b_Q$) is indirect, by changing $\alpha_{t,s}$ (routing/selection) and is therefore not represented as a simple additive term in $u_{\text{bias}}^{(\ell)}$. The key bias contribution ($b_K$) cancels in softmax normalization (standard dot-product attention) and therefore does not contribute to $\alpha_{t,s}$ or to the head output via this route.

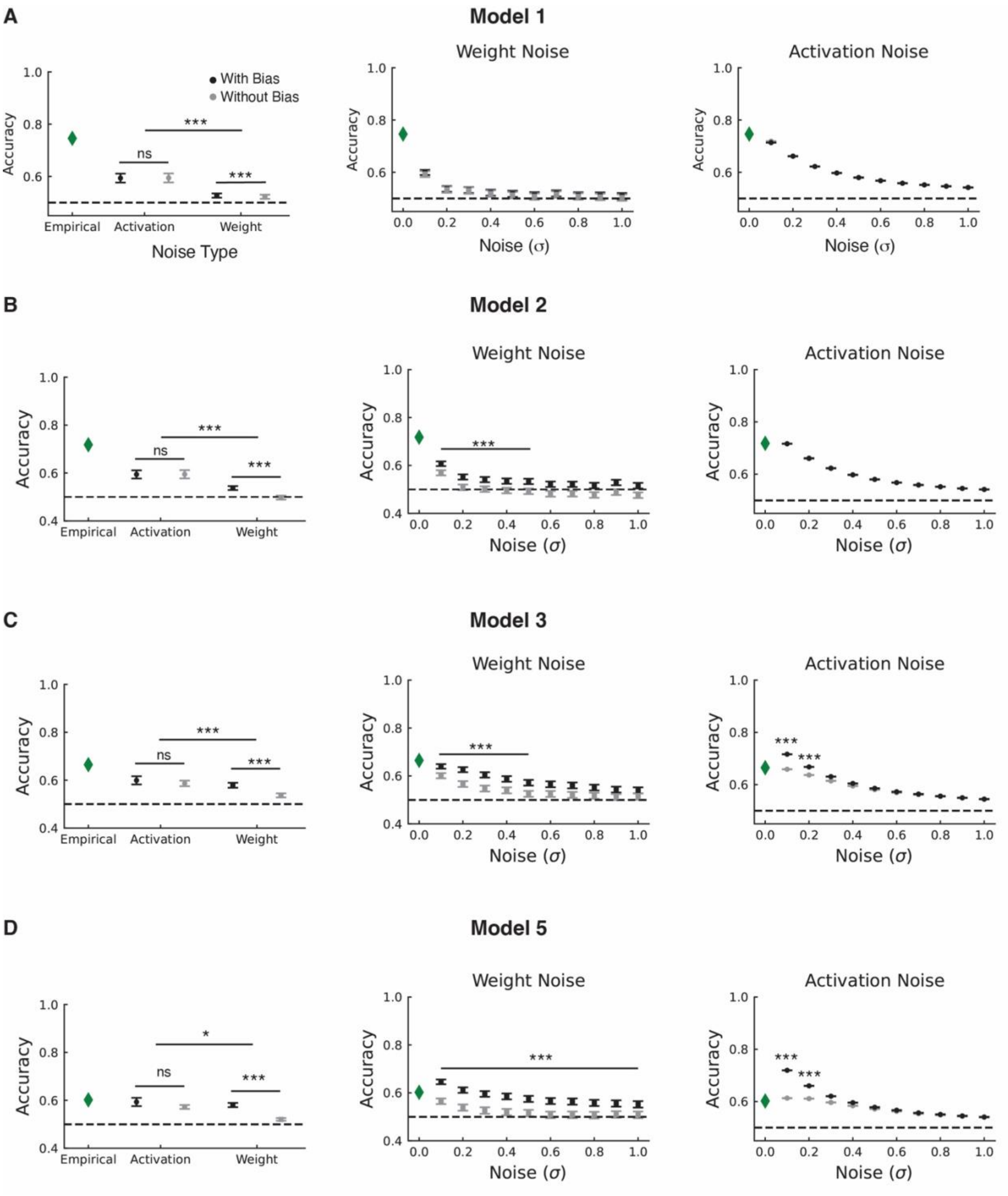


**Figure S1. Bias component preserves performance against noise in the weights.** (A) Model 1. Average classification accuracy after adding noise (*left*). Classification accuracy for each noise level with added noise to the weights (*middle*) or activations (*left*). Green diamond represents the empirical performance with no added noise ( $\sigma = 0$) is shown for reference. Black and gray dots represent the average classification value across all noise levels for with and without bias term in the classification, respectively. The dashed line indicates chance performance (0.5). Error bars show mean ± sem. (B) Same as panel A, but for Model 2. (C) Same as panel A, but for Model 3. (D) Same as panel A, but for Model 5. Statistical significance was determined using paired-sample *t*-test and FDR corrected. ***$P < 0.001$; ns = not significant.

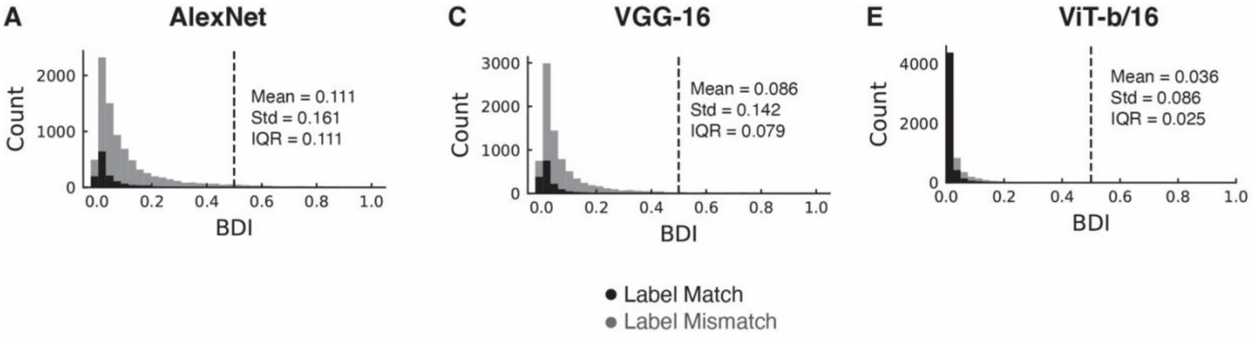


**Figure S2. Distribution of BDI values.** (A) AlexNet. (B) VGG-16. (C) ViT-b/16. Analysis is based on 10,000 images from the validation set part of Tiny-ImageNet-1K.

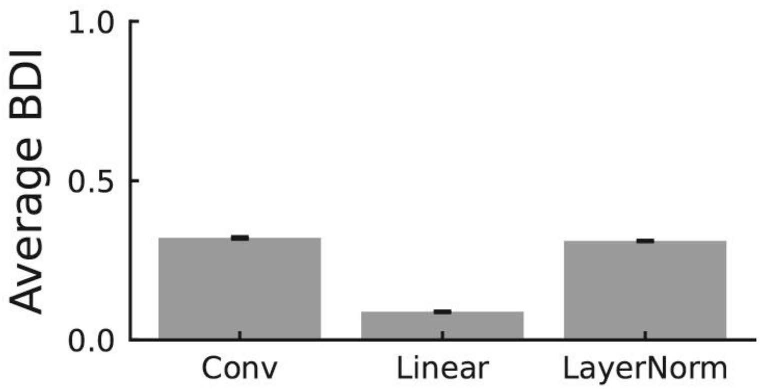


**Figure S3. ViT-b/16 average BDI across layer groups.** BDI is estimated by averaging the BDI values across Linear and LayerNorms, respectively. Note that that there is only one convolutional layer in ViT-b/16.

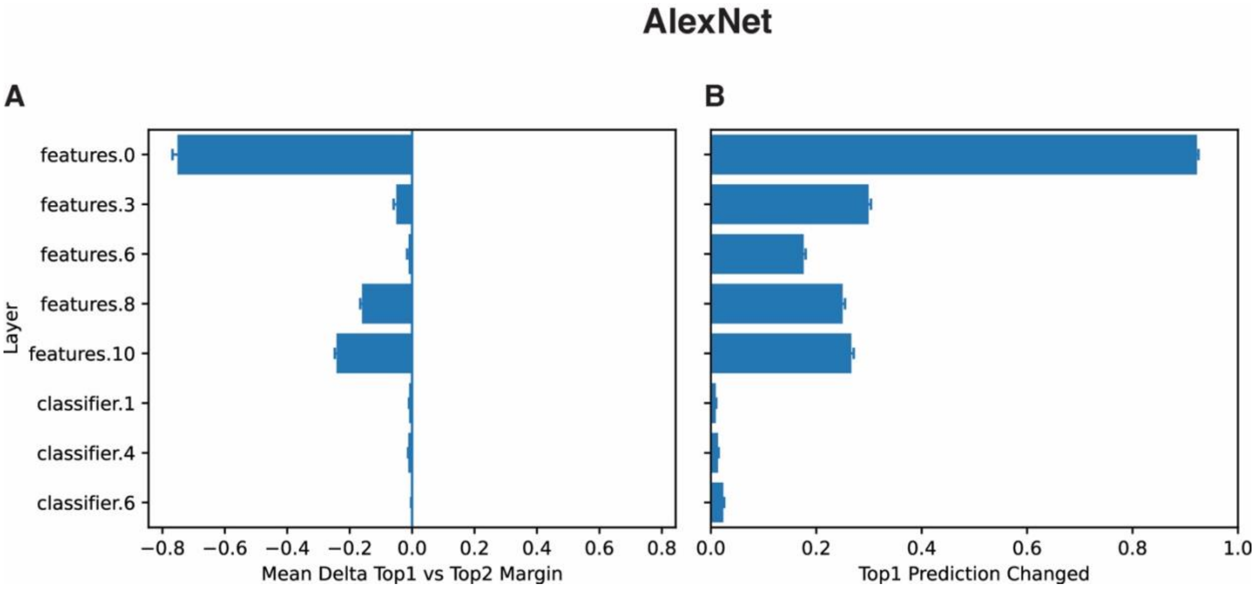


**Figure S4. Clamping the bias modulated the decision margin and top1 predicted class in AlexNet.** (A) The extent to which the margin changed after clamping the bias to zeros. (B) The proportion of top-1 predictions that changes between clamped and unclamped bias. The bias was clamped to zero for one layer at a time.

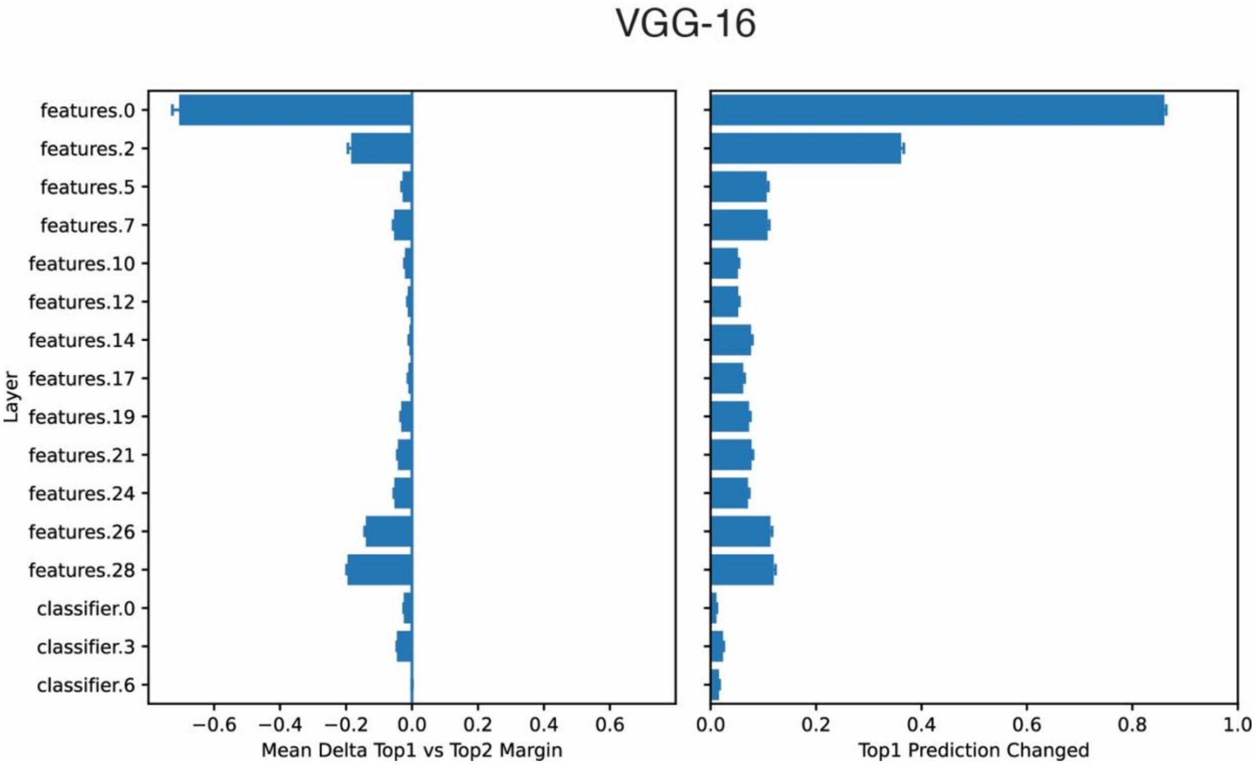


**Figure S5. Clamping the bias modulated the decision margin and top1 predicted class in VGG-16.** (A) extent to which the margin changed after clamping the bias to zeros. (B) The proportion of top-1 predictions that changes between clamped and unclamped bias. The bias was clamped to zero for one layer at a time.

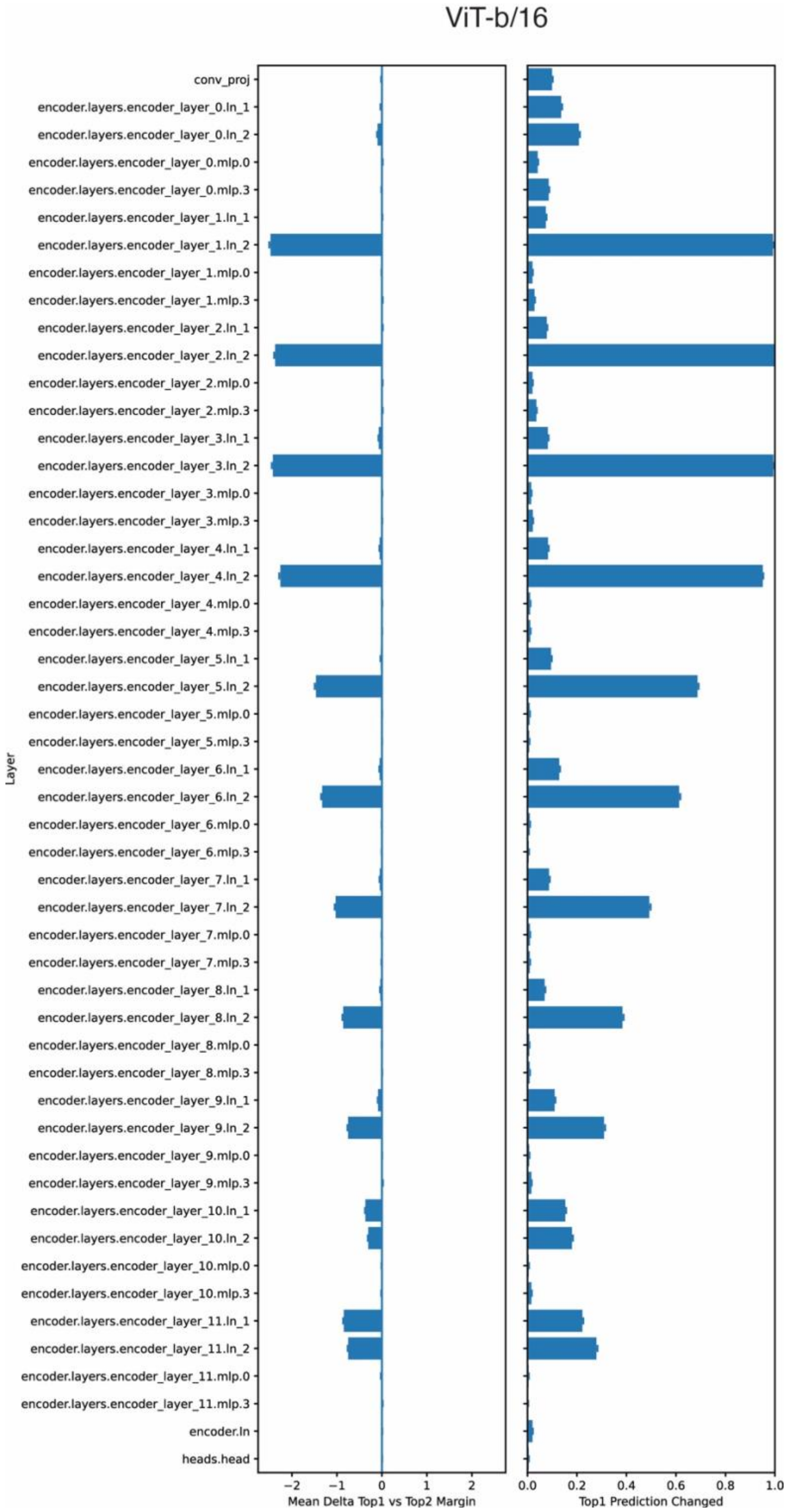


**Figure S6. Clamping the bias modulated the decision margin and top1 predicted class in ViT-b/15.** (A) The extent to which the margin changed after clamping the bias to zeros. (B) the proportion of top-1 predictions that changes between clamped and unclamped bias. The bias was clamped to zero for one layer at a time.

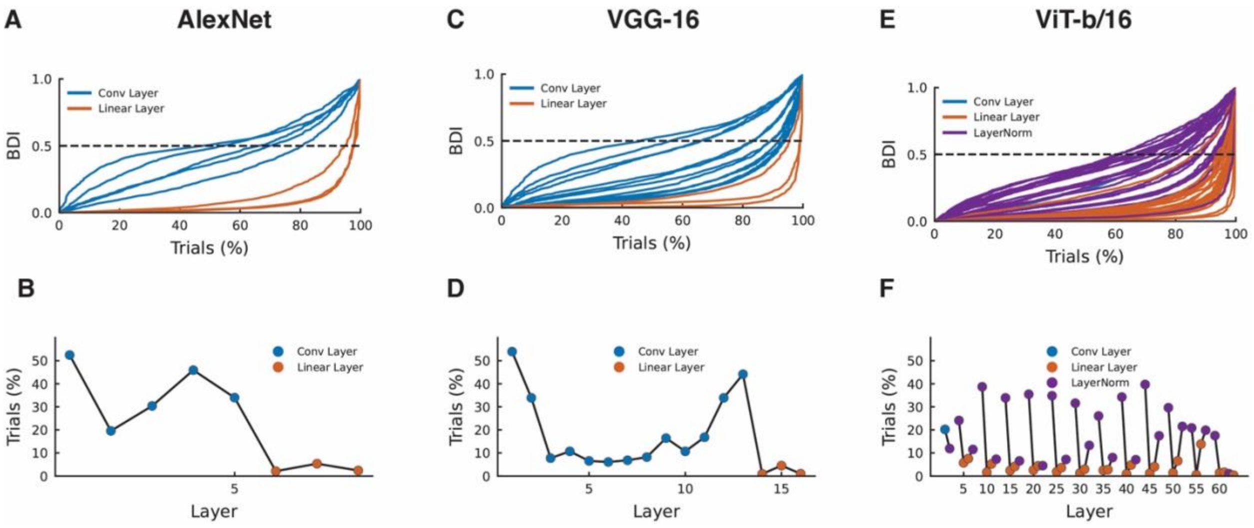


**Figure S7. Layer-resolved bias dominance evaluated on ImageNet.** (A) Layer-resolved BDI profile for AlexNet. Colored lines correspond to layers spanning convolutional and fully connected layers that contain a bias component. Analysis is based on 1,000 images from the ImageNet dataset. Trials are ranked by BDI values. The dashed line indicates the bias-dominance criterion BDI = 0.5. (B) Percentage of bias-dominant trials with BDI > 0.5 across layers for AlexNet. Layers are ordered sequentially according to their execution order on the input, and only layers with bias components are included. (C) VGG-16 results shown as in panel A. (D) Percentage of bias-dominant trials with BDI > 0.5 across layers for VGG-16. (E) ViT-b/16 results shown as in panel A. (F) Percentage of bias-dominant trials with BDI > 0.5 across layers for ViT-b/16.

s

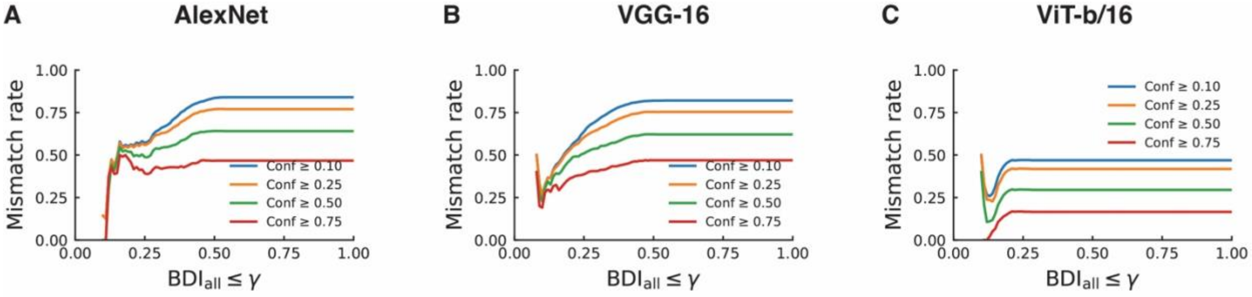


**Figure S8. Relationship between mismatch rate and BDI threshold at fixed confidence levels.** Panels show AlexNet (A), VGG-16 (B), and ViT-b/16 (C). For each model, trials were grouped by minimum confidence threshold, and the mismatch rate was plotted as a function of $BDI_{all} \leq \gamma$.

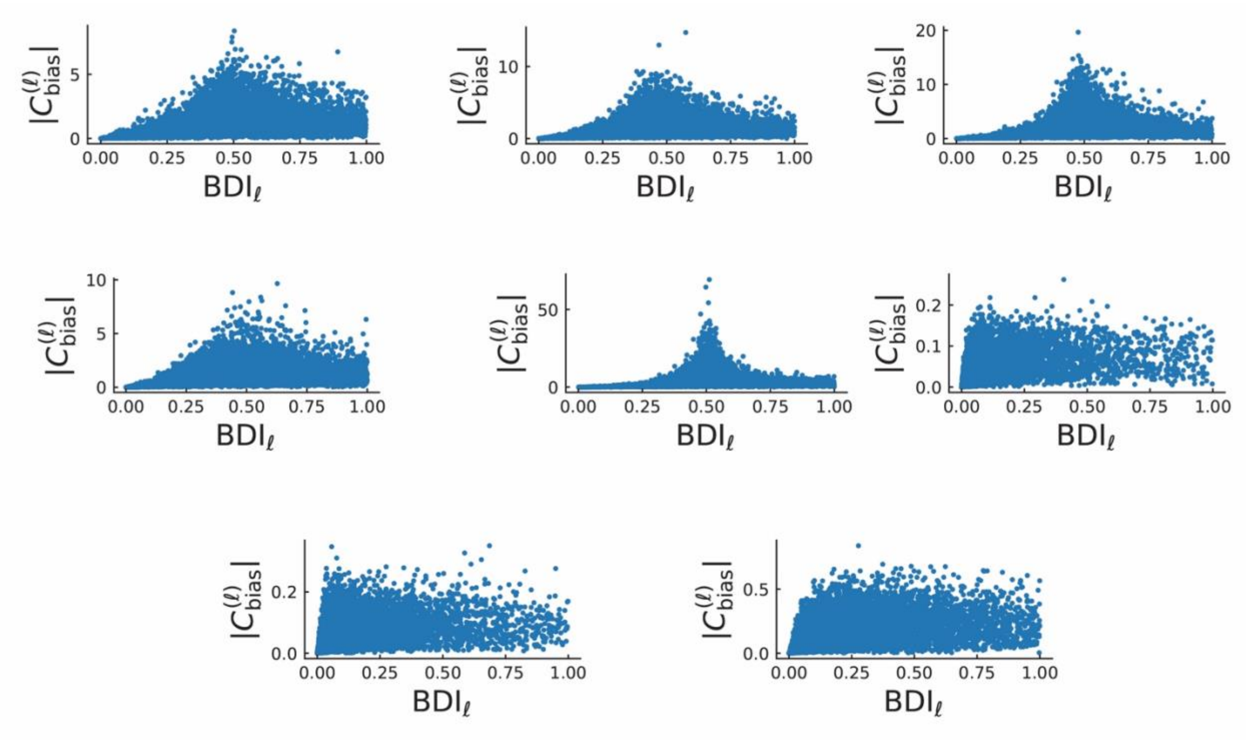


**Figure S9. AlexNet relationship between BDI and bias magnitude.** For AlexNet, BDI was positively associated with $|\ C_{\text{bias}}^{(\ell)}\ |$ the relationship was far from deterministic, with substantial dispersion in BDI values at fixed $|\ C_{\text{bias}}^{(\ell)}\ |$.

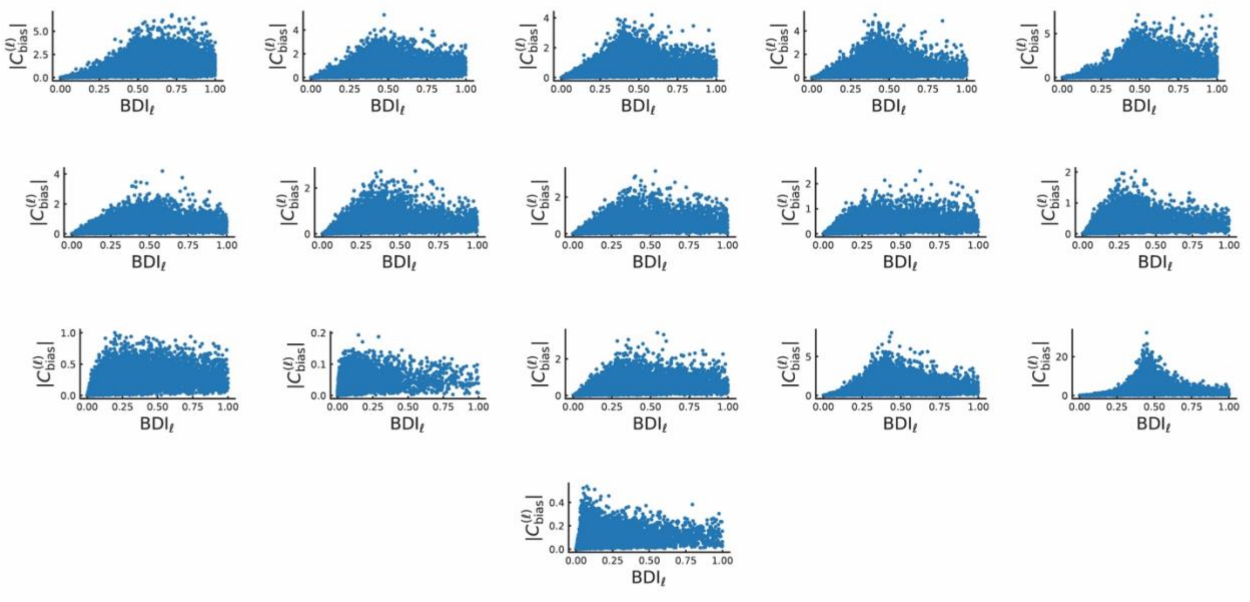


**Figure S10. VGG-16 Relationship between BDI and bias magnitude.** For VGG-16, the relationship between BDI and | $C_{\text{bias}}^{(\ell)}$ | exhibited substantial dispersion in BDI values at fixed | $C_{\text{bias}}^{(\ell)}$ |.

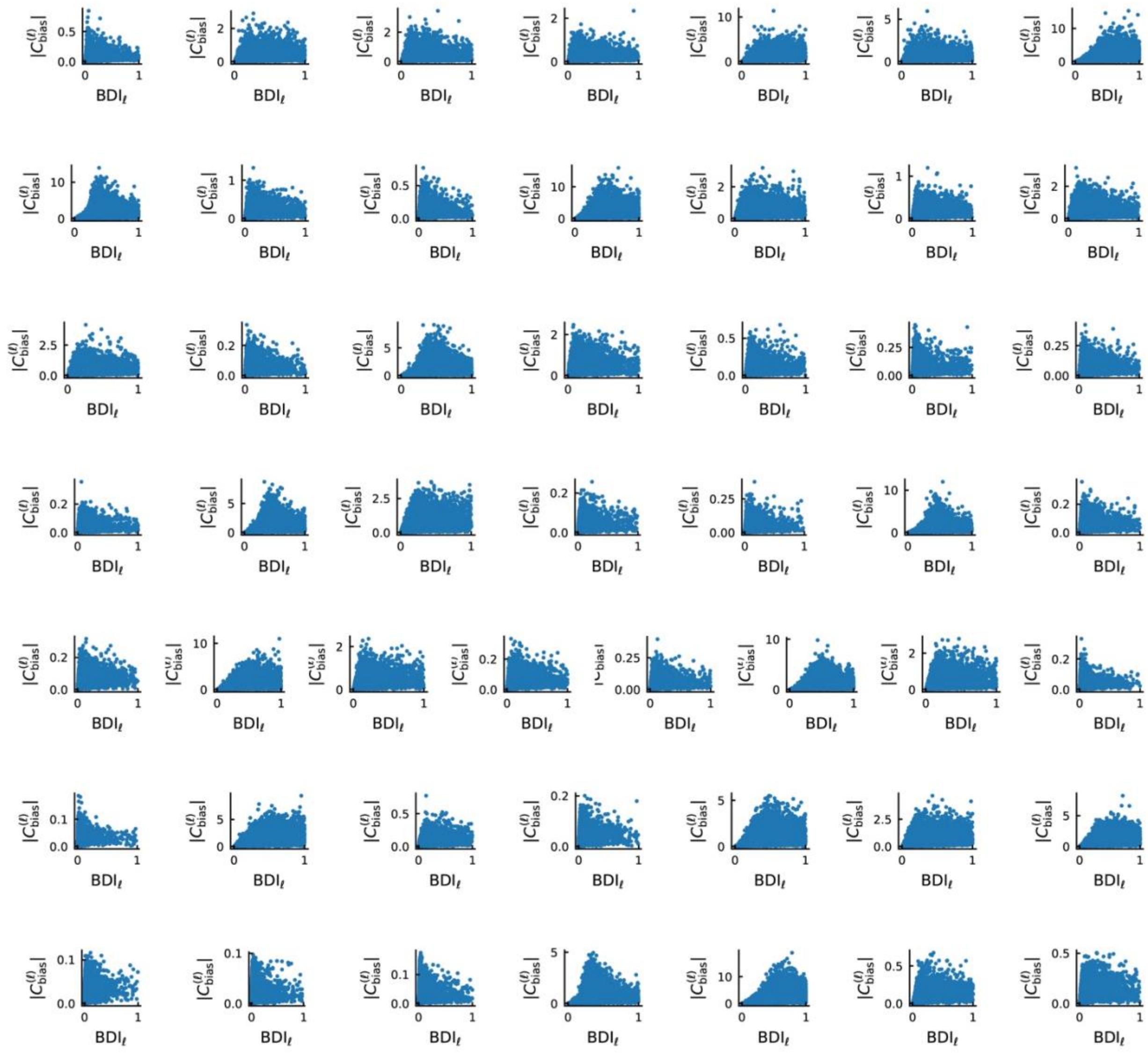


**Figure S11. ViT-b/16 Relationship between BDI and bias magnitude.** For ViT-b/16, the relationship between BDI and | $C_{\text{bias}}^{(\ell)}$ | exhibited substantial dispersion in BDI values at fixed | $C_{\text{bias}}^{(\ell)}$ |.

### Table S1: BERT Completed Prompts

| Prompt | Top1 Token | Top2 Token | Top1 Confidence | BDI Max |
|---|---|---|---|---|
| She reported months of polyuria and polydipsia; HbA1c returned at 11.2%, and the clinician suspected [MASK]. | infection | cancer | 0.0812 | 0.8390 |
| On arrival the patient was febrile and tachycardic; blood cultures later grew gram-positive cocci, consistent with [MASK]. | symptoms | this | 0.2366 | 0.9971 |
| The ECG showed ST elevation in leads II, III, and aVF; they treated it as an acute [MASK]. | phase | condition | 0.0797 | 0.9145 |
| CT angiography demonstrated a filling defect in the right pulmonary artery, most consistent with [MASK]. | age | mri | 0.0802 | 0.9229 |
| The chest radiograph showed a lobar consolidation; given the productive cough, the leading concern was [MASK]. | pneumonia | health | 0.1614 | 0.9841 |
| After starting the new antibiotic, the patient developed diffuse hives and wheezing, suggesting an [MASK]. | infection | ##emia | 0.4403 | 0.9820 |
| The biopsy showed caseating granulomas and acid-fast bacilli, raising concern for [MASK]. | cancer | him | 0.0872 | 0.8190 |
| He presented with unilateral facial droop and slurred speech; the team activated a [MASK] alert. | red | yellow | 0.2024 | 0.9790 |
| Lumbar puncture showed elevated opening pressure and neutrophilic pleocytosis, supporting [MASK]. | diagnosis | survival | 0.1055 | 0.9903 |
| MRI revealed periventricular demyelinating lesions, and the neurologist discussed [MASK] as the likely diagnosis. | this | it | 0.215 | 0.9895 |
| Given the persistent wheeze, nocturnal symptoms, and reversible obstruction on spirometry, they initiated an [MASK] inhaler. | emergency | asthma | 0.4132 | 0.9846 |
| Because the INR was supratherapeutic and there was active bleeding, they administered [MASK]. | it | antibiotics | 0.1996 | 0.9667 |
| With rising creatinine and oliguria after contrast exposure, the note favored contrast-induced [MASK]. | blindness | inflammation | 0.0526 | 0.9852 |
| Because the pain radiated to the back and lipase was markedly elevated, the working diagnosis was [MASK]. | correct | made | 0.0647 | 0.9711 |
| After a witnessed tonic-clonic event with post-ictal confusion, the chart documented a [MASK]. | econstructior | recovery | 0.029 | 0.9911 |
| Due to the new systolic murmur and positive blood cultures, the team evaluated for [MASK]. | infection | treatment | 0.0632 | 0.9485 |
| A week after immobilization, the calf became swollen and tender; ultrasound confirmed [MASK]. | this | it | 0.6669 | 0.9458 |
| Because the sodium was 119 mmol/L with low serum osmolality, they managed severe [MASK]. | symptoms | bleeding | 0.096 | 0.8462 |
| With progressive dyspnea and bilateral crackles plus elevated BNP, they treated for [MASK]. | depression | ms | 0.0814 | 0.9588 |
| Given episodic palpitations and an irregularly irregular rhythm, the diagnosis was [MASK]. | invalid | confirmed | 0.1941 | 0.9758 |
| Although the CT was negative for hemorrhage, the sudden severe headache kept [MASK] on the differential. | working | insisting | 0.2637 | 0.8806 |
| The report noted no evidence of [MASK], but recommended follow-up imaging given the borderline findings. | infection | disease | 0.2875 | 0.9905 |
| Despite normal troponins, the clinical picture remained concerning for [MASK]. | decades | years | 0.1111 | 0.9518 |
| The clinician questioned whether the symptoms reflected anxiety rather than [MASK]. | depression | fear | 0.5296 | 0.9891 |
| The note favored viral illness over bacterial [MASK] given the negative procalcitonin. | infection | infections | 0.4297 | 0.9329 |
| Assessment: intermittent claudication with diminished distal pulses; impression most consistent with peripheral [MASK]. | pulses | pulse | 0.3788 | 0.9753 |
| Impression: enlarged ventricles with transependymal flow; findings suggest [MASK]. | ##ive | otherwise | 0.1041 | 0.9514 |
| Summary: microcytic anemia with low ferritin; pattern consistent with [MASK]. | age | symptoms | 0.1611 | 0.9809 |
| Impression: elevated TSH and low free T4; consistent with primary [MASK]. | stress | development | 0.0388 | 0.8750 |
| Assessment: tremor, rigidity, and bradykinesia; features compatible with [MASK]. | surgery | mri | 0.0939 | 0.9317 |
| Given the repeated missed deadlines and inconsistent explanations, the manager concluded the issue was [MASK]. | resolved | serious | 0.1697 | 0.9847 |
| Although the sky was clear earlier, the sudden gusts and dark clouds suggested an approaching [MASK]. | storm | typhoon | 0.8124 | 0.9844 |
| After reviewing the receipts and bank statements, the auditor suspected [MASK]. | fraud | corruption | 0.4159 | 0.9540 |
| Because the device overheated only under load and the fan never spun up, the likely failure was the [MASK]. | same | fan | 0.1752 | 0.9030 |
| The committee agreed that the unexpected results were driven by a hidden [MASK] in the dataset. | flaw | vulnerability | 0.2034 | 0.9455 |
| The solution arrived not as inspiration, but as careful [MASK]. | consideratior | practice | 0.1507 | 0.9563 |
| On the moor, the fog erased distance and replaced it with [MASK]. | water | snow | 0.1384 | 0.9653 |
| The howl carried over the marsh, turning skepticism into [MASK]. | fear | anger | 0.1943 | 0.8018 |
| He studied the legend and found its power lay in cultivated [MASK]. | land | plants | 0.0990 | 0.9159 |
| A lantern flickered in the ruins, and the silence became [MASK]. | awkward | oppressive | 0.0964 | 0.9984 |
| He stared at what he had made and felt triumph transform into [MASK]. | fear | joy | 0.1072 | 0.8289 |
| The creature,Äôs quiet request exposed the thinness of his [MASK]. | skin | throat | 0.1644 | 0.9083 |
| In the mountain pass, the confession sounded less like guilt than [MASK]. | fear | anger | 0.0998 | 0.9661 |
| He realized too late that pursuit can become a kind of [MASK]. | obsession | habit | 0.1631 | 0.9188 |
| The polite fa√ßade held, but the handwriting betrayed [MASK]. | it | him | 0.5046 | 0.9887 |
| At the door, the servant,Äôs hesitation spoke of [MASK] more than certainty. | something | fear | 0.2977 | 0.9917 |
| He tried to reason it away, yet the evidence insisted on [MASK]. | it | silence | 0.1382 | 0.9597 |
| The final letter explained everything, and still left him with [MASK]. | doubts | nothing | 0.1502 | 0.8354 |
| The compliment lingered like perfume, masking a sharper [MASK]. | edge | sting | 0.3201 | 0.9721 |

#### Table S1 Continued:

| Prompt | Top1 Token | Top2 Token | Top1 Confidence | BDI Max |
|---|---|---|---|---|
| He laughed easily, but the mirror returned only [MASK]. | briefly | slightly | 0.1723 | 0.9777 |
| The room remained beautiful, while his thoughts darkened into [MASK]. | nothing | darkness | 0.1133 | 0.9153 |
| He sought distraction, yet every pleasure carried a trace of [MASK]. | pain | fear | 0.0958 | 0.9394 |
| At the window, the pale mark on her throat turned fear into [MASK]. | anger | hope | 0.1055 | 0.8548 |
| The journal entries grew stranger, threaded with mounting [MASK]. | anxiety | tension | 0.1271 | 0.9700 |
| They agreed to keep watch, though each hour deepened [MASK]. | dusk | darker | 0.1080 | 0.8412 |
| In the carriage at night, the countryside felt ruled by [MASK]. | darkness | nature | 0.0965 | 0.8579 |
| He described the distant future, and the listeners traded amusement for [MASK]. | amusement | sympathy | 0.1031 | 0.9486 |
| The silent ruins suggested not progress, but slow [MASK]. | ##ness | progress | 0.2292 | 0.9591 |
| He chased an answer across centuries and found only [MASK]. | one | truth | 0.6123 | 0.8473 |
| When he returned, he spoke softly, as if afraid of [MASK]. | something | me | 0.6818 | 0.9946 |
| The first cylinder opened, and curiosity gave way to [MASK]. | fear | excitement | 0.1275 | 0.9553 |
| The streets emptied faster than belief, driven by [MASK]. | traffic | death | 0.1268 | 0.9962 |
| He heard the distant machinery and recognized the sound of [MASK]. | it | footsteps | 0.0594 | 0.9500 |
| Even the calm voice on the radio cracked into [MASK]. | silence | laughter | 0.2807 | 0.9745 |
| The conversation bent in impossible loops, and she answered with [MASK] to keep up. | nothing | effort | 0.1501 | 0.9816 |
| The rules changed mid-sentence, producing instant [MASK] among the guests. | laughter | applause | 0.0976 | 0.9821 |
| A grin appeared without a body, and she felt a tug of [MASK]. | jealousy | fear | 0.0869 | 0.9058 |
| The tea never ended, and the politeness became [MASK]. | stronger | awkward | 0.0390 | 0.9438 |
| The path looked simple, yet each step demanded [MASK]. | attention | caution | 0.0991 | 0.9466 |
| The great voice boomed, but behind it she sensed [MASK]. | it | danger | 0.1286 | 0.9217 |
| The companions argued, and the dispute softened into [MASK] when the road narrowed. | one | confusion | 0.1532 | 0.9722 |
| When the curtain shifted, fear turned quickly into [MASK]. | anger | excitement | 0.2000 | 0.9645 |
| The cold taught him a brutal kind of [MASK]. | lesson | loyalty | 0.1054 | 0.9604 |
| He felt the old instincts rise, stronger than training, like [MASK]. | instinct | magic | 0.0617 | 0.9077 |
| The campfire stories sounded distant, drowned by [MASK]. | noise | voices | 0.1070 | 0.9956 |
| The final choice arrived as quiet [MASK], not a shout. | acceptance | silence | 0.0839 | 0.9809 |
| The law of the jungle was recited calmly, but it demanded [MASK]. | respect | obedience | 0.1258 | 0.9332 |
| He listened to the council and learned that belonging requires [MASK]. | respect | trust | 0.1071 | 0.9619 |
| The warning was gentle, yet it carried real [MASK]. | meaning | fear | 0.4415 | 0.9576 |
| In the night, the forest seemed to breathe [MASK]. | heavily | again | 0.1934 | 0.9966 |
| The crowd‚Äôs attention fixed on her, and shame hardened into [MASK]. | anger | fury | 0.3603 | 0.8381 |
| The minister spoke with elegance, but the silence answered with [MASK]. | silence | respect | 0.0565 | 0.9426 |
| The token was meant to punish, yet it became [MASK] in her hands. | useless | stuck | 0.1995 | 0.9728 |
| When the truth threatened to surface, the town chose [MASK]. | him | it | 0.1766 | 0.9030 |
| He watched the captain‚Äôs gaze and saw obsession sharpen into [MASK]. | steel | anger | 0.1444 | 0.9952 |
| The sea was calm, yet the crew moved with [MASK] at the thought of the hunt. | excitement | anticipation | 0.1914 | 0.9495 |
| The sermon lingered, turning the voyage into [MASK]. | madness | another | 0.0412 | 0.9772 |
| A glimpse of the whale transformed awe into [MASK]. | fear | horror | 0.1625 | 0.9504 |
| In the cell, he learned patience as a form of [MASK]. | punishment | torture | 0.2156 | 0.9079 |
| The disguise fit perfectly, but it carried the weight of [MASK]. | it | death | 0.0512 | 0.9804 |
| He smiled at his enemies, practicing [MASK] with flawless manners. | them | himself | 0.6725 | 0.9365 |
| The plan unfolded slowly, each step powered by [MASK]. | adrenaline | energy | 0.0895 | 0.9029 |
| The duel was postponed, not cancelled, and the men replaced anger with [MASK]. | fear | anger | 0.0769 | 0.9678 |
| A letter arrived sealed tight, promising both favor and [MASK]. | friendship | forgiveness | 0.0972 | 0.9945 |
| He boasted loudly, hiding a flicker of [MASK]. | amusement | humor | 0.1834 | 0.9838 |
| The oath was sworn, and suddenly everything felt like [MASK]. | glass | magic | 0.0589 | 0.9882 |
| He offered to trade chores, disguising his scheme as [MASK]. | his | business | 0.0706 | 0.9128 |
| The town‚Äôs excitement grew, fueled by rumor and [MASK]. | gossip | rumor | 0.1966 | 0.9807 |
| They crept into the graveyard, and bravado slipped into [MASK]. | place | him | 0.1393 | 0.9395 |
| The treasure talk sounded like play, until it turned into [MASK]. | laughter | music | 0.2066 | 0.9684 |
| The night ride began as pride and ended in [MASK]. | victory | death | 0.0926 | 0.9971 |
